# Rapid Intracellular Delivery of Human Heat Shock Protein 72 Inhibits Neurodegeneration and Oxidative Damage After a Traumatic Brain Injury

**DOI:** 10.64898/2026.05.03.722564

**Authors:** Allen Chan, Manda Saraswati, Krutik Patel, Sonia Su, Anna Su, Peethambaran Arun, Peyton Politewicz, Joni Ricks-Oddie, Dallas Hack, Robert Nishimura, Stephen T. Hobson, Richard A. Richieri, Karolina Krasinska, Courtney L. Robertson, Missag H. Parseghian

## Abstract

Fv-HSP72 is a rapid cell-penetrating human heat shock protein for the treatment of traumatic organ injuries. We have shown this re-engineered protein (HSP72) is capable of crossing the blood brain barrier (BBB) of rats suffering a controlled cortical impact (CCI) and remains in brain tissue for up to 12 hours; long after clearance from the cortex of uninjured rats. Peptide sequences unique to Fv-HSP72 allow for its differential detection from endogenous HSP72. Male Sprague-Dawley rats were divided into 10 groups of n=10 with those animals receiving a CCI subjected to a unilateral cortical contusion simulating a moderate to severe brain injury using an electronically controlled pneumatic impact device. Control groups were either uninjured (Sham), injured (TBI Only), or injured and given buffer (TBI+Vehicle). Rats treated with one of three Fv-HSP72 variants were dosed at 10 or 30mg/kg 15m post-impact, then sacrificed 48 hours later. Cortical tissues were extracted from the ipsilateral and contralateral hemispheres for biomarker analysis. Here we report results of our drug inhibiting neurodegeneration based on five biomarkers (NF-L, pNF-H, pTau [T181, T231, S396]). These results were statistically significant, especially for one of the Fv-HSP72 variants, when comparing differences both between treatment groups and within groups (i.e. when comparing ipsi-vs. contralateral hemispheres). Significant inhibition of oxidative stress (3-NT) and inflammatory (IL-6) biomarkers were also observed (both p<0.0001). With similar results obtained for a blast injury model being published elsewhere, the analyses suggest Fv-HSP72 is neuroprotective following a direct impact brain injury.

**One sentence summary:** This study describes the effectiveness of a biologic agent, Fv-HSP72, in significantly inhibiting neuronal tissue damage in the brain when administered after a direct cortical impact.

---

It is estimated 69 million individuals worldwide sustain a traumatic brain injury (TBI) each year^1^. TBI can lead to neurodegeneration, oxidative stress, inflammation, and disruption of the blood brain barrier (BBB). In response, a variety of cells in the brain express heat shock protein 72 (HSP72) to counter the ensuing apoptotic damage. Using a controlled cortical impact (CCI) model, researchers have shown that these symptoms can be attenuated with the upregulation of HSP72 (HSPA1A in the current literature, and historically known as HSP70 or HSP70i)^2,3^. In two independent laboratories, HSP72^-/-^ knockout (KO) mice had increased lesion sizes, brain hemorrhage and worsened behavioral outcomes compared to wildtype (Wt) mice^2^. The authors found increased expression of matrix metalloproteinases (MMPs) -2 and -9 after injury, and suggested it led to further breakdown of the extracellular matrix and the BBB, hence the brain edema and hemorrhage characteristic of TBI. The other lab found inhibition of HSP70 and HSP110 resulted in increased expression of pro-inflammatory and oxidative stress genes after TBI. Furthermore, they demonstrated that drugs that increase the level of HSP70/HSP110 reduced apoptosis, inflammatory cell infiltration and gliosis after TBI, and improved behavioral outcomes^3^. Similarly, transgenic mice over-expressing HSP72 had smaller brain lesion sizes, decreased hemorrhage, better behavioral outcomes and reduced expression of MMPs-2 and -9 in the same CCI model^2^.

Natural HSP72 induction can take several hours in the central nervous system (CNS)^4^, up to 6 hours in the case of ischemia^5^, which is a lag time that may be detrimental to a TBI patient needing immediate intervention. Rapid delivery of HSP72 to brain tissue protects cells from injury^6^ and a solution already exists that avoids the typical delays necessary for induction of the HSP72 gene. To prevent neurological damage resulting from TBI, we exogenously deliver a re-engineered HSP72 into neuronal tissues and their supporting vasculature. This delivery is accomplished as HSP72 is fused with the humanized scFv fragment of an antibody, mAb 3E10, that binds extracellular DNA and is being developed as an intracellular transporter for protein therapeutics (Fv-HSP72). 3E10 penetration occurs through a specific nucleoside salvage channel found in most cells^7,8^, known as the equilibrative nucleoside transporter 2 (ENT2; *SLC29A2*) whose expression is relatively high in brain tissue^9,10^. DNA fragments are abundant and accessible during cell necrosis after tissue injury^11,12^. 3E10 penetration of non-dividing primary rat cortical neurons has been shown *in vitro*^13^ and Fv-HSP72 has demonstrated *in vivo* efficacy in cerebral^6^ and cardiovascular^14^ infarction models. Selectivity of Fv-HSP72 targeting *in vivo* occurs because tissues undergoing significant cell injury possess a high concentration of extracellular DNA. Salvaging of the DNA by surrounding cells, through the ENT2 channel, provides 3E10 fused to HSP72 the opportunity to enter those cells and inhibit further cell death.

## Methods

### Materials

Fv-HSP72 is the name given to a class of fusion proteins being developed by Rubicon Biotechnology (Irvine, CA). Six humanized variants of the 3E10 scFv fragment have been fused to several modified forms of human HSP72 that eliminate sites of glycosylation, oxidation and acid-labile cleavage. The Fv-HSP72s are ∼98kD proteins synthesized in CHO cells, purified by chromatography, and formulated to 6mg/mL in PBS with 5% sucrose. For this study, two humanized 3E10 variants were chosen, codenamed RBB010 and RBB012, based on *in vitro* DNA binding kinetics and neural cell uptake (data not shown). Fusion of the 3E10 subunit to the HSP72 occurs with one of two cleavable amino acid linkers: a “Swivel” linker (RBB010, RBB012) or a cathepsin B cleavable one (RBB012-CTB). Product potency is tested both in the 3E10 subunit, which binds the target, and the HSP72 subunit, which requires a functional ATP subdomain (data not shown).

All animal studies were conducted with male Sprague-Dawley rats (CD-strain; Charles River), weighing 225-350g, at Johns Hopkins Univ. (JHU; Baltimore) after review and approval by JHU’s IACUC and US Army’s ACURO teams. The experiments reported herein were conducted in compliance with the Animal Welfare Act and per the principles set forth in the “Guide for Care and Use of Laboratory Animals,” Institute of Laboratory Animal Resources, National Research Council, National Academy Press, 2011.

### CCI Injury

Those animals receiving a TBI were subjected to a unilateral cortical contusion using an electronically controlled pneumatic impact device as previously described^15^. All animals are anesthetized with isoflurane (2-2.5%) and placed in a stereotaxic frame, the skin is retracted and a 6 mm unilateral craniotomy is performed that is centered between the bregma and lambda. The skull cap is removed without disruption of the underlying dura. The exposed brain is injured using a 6 mm tip that impacts the cortex at 5.5 m/s to a depth of 2mm with a duration of 50msec. For these studies, the left hemisphere of the brain was the side that received the contusion.

Following injury, the bone flap was replaced, the craniotomy sealed with an acrylic mixture (Koldmount, Albany, NY), and the scalp incision was closed with interrupted sutures. At the completion of surgery, isoflurane was discontinued, and rats were awakened and returned to their cages. Sham rats (blank implanted) underwent identical surgeries, with the exclusion of the CCI.

### Pharmacokinetics (PK) using Liquid Chromatography (LC) - Mass Spectrometry (MS)

Rats received a **10mg/kg** dose of Fv-HSP72 intravenously (IV) by tail vein injection 15m post-CCI. Naïve rats were put under isoflurane anesthesia and injected with the drug without any other surgical procedures. Naïve rats received a 10mg/kg dose of Fv-HSP72 IV by tail vein injection 15m post-anesthesia. Both, naïve rats and those exposed to a CCI were divided into 3 groups (n=5/group) with one group sacrificed 1h after drug injection, and two other groups at 4h and 12h post-injection. All 3 Fv-HSP72s were subjected to PK analysis of the brain and plasma tissues, making for a total of 18 groups.

#### Tissue Extraction for PK Analysis

Brain and plasma were harvested from rats 1, 4, and 12 hrs post-injection. The cortical tissues required further processing and homogenization prior to LC-MS with brain tissues harvested at the ipsilateral and contralateral hemispheres separately for analysis. In brief, solid cortical tissue samples were washed with PBS prior to removal from the rat, they were weighed and then tissues were frozen on dry ice and shipped to Stanford for further processing. Brain samples were homogenized in 1mL of PBS, with the addition of HALT protease inhibitors (Thermo), using Lysing Matrix D beads (MP Biomedicals) and a bead beater (Benchmark Scientific D2400-R 24) run for 30s at 3m/s before putting the extract back on ice to avoid sample overheating. The bead beating and cooling sequence was repeated 3 times. The protein content in the homogenate was determined using a Bradford assay to normalize the results from the samples.

#### Identifying Suitable Peptide Fragments for PK Analysis

Proteomic analysis of our Fv-HSP72 variants was conducted in a QEHFx mass spectrometer (Thermo) coupled with an I-class UPLC chromatography system (Waters). Briefly, the purified Fv-HSP72 variants in a 5% sucrose/PBS solution were subjected to reduction (with DTT) and alkylation (acrylamide) followed by proteolytic digestion with Trypsin/Lys-C. After the C18 cleanup step the peptide mixture was analyzed via LC-MS/MS. The data dependent acquisition (DDA) mode and subsequent data base search resulted in detection of 11,000-12,000 peptides representing Fv-HSP72 protein. Several unique peptides representing various parts of the fusion protein were chosen from the proteomic analysis. The selected peptides were chemically synthetized by Anaspec to be used for the absolute quantitative assay setup on a triple quadrupole LC-MS/MS system. The commercially obtained **<u>unlabeled peptide</u>** standards were used for LC-MS/MS parameter optimization and calibration curve characterization; while, **<u>stable</u> <u>isotope “labeled”</u>** versions of the same peptides were used for matrix effect determination and monitoring, as well as normalization of results. None of the chosen peptides appear to form albumin-drug complexes, a serious limitation for quantitation of proteins in blood using MS.

#### The PK Assay

The absolute quantitation of selected peptides representing the target protein was set up on a Xevo TQ-XS triple quadrupole MS coupled with an Acquity I-class UPLC system and operated in positive electrospray ionization mode. Each peptide was monitored via selected reaction monitoring (SRM) where unique pairs of precursor and fragment ions, representing each target peptide, are detected. There were 2-3 SRM transitions selected for each peptide, both unlabeled and labeled.

Chromatographic separation was performed on an Acquity UPLC BEH C18 column (2.1 x 100 mm, 1.7 μm) with a gradient elution from 5% to 98% B (10m total run time), where mobile phase A was 0.2% formic acid in water and mobile phase B was 0.2% formic acid in acetonitrile. The flow rate was 0.3 mL/min and column temperature was 35°C. Sample injection volume was 5µl.

#### Sample Analysis

Every sample was injected twice to obtain 2 technical repeats (as opposed to “biological repeats” which are obtained from multiple animals being treated the same way). If the Signal to Noise (S/N) ratio was ≥10:1 for the two technical repeats, then the values obtained were considered reliable. Values <10:1 are often detectable, but not reliably quantifiable.

#### Calculating pg/mg from fmol/µL for the brain tissue samples

While the results obtained from the mass spec are in fmol/µL, they had to be converted into pg of ITP peptide per milligrams of brain tissue in order to normalize the results across all of the samples. For the analysis, Stanford took 100µL of each homogenate, processed and reconstituted it in 100µL of sample buffer and injected 5µL into the LC-MS/MS system. The fmol/µL value obtained corresponds to the concentration of each biomarker peptide in the tissue homogenate. The fmol/µL were converted to ng/µL by multiplying with the molecular weight of the particular peptide. The ng/µL were normalized by dividing in milligrams with the starting brain tissue mass for each sample. Since the concentrations were low, they were converted from ng into picograms of Fv-HSP72 protein/mg of brain tissue. These are the values plotted in **Figure 10**.

### Screening Fv-HSP72 Variant Efficacy in the CCI Model

#### Study Groups

Rats were divided into 10 groups (n=8/group). The Sham control group underwent the anesthesia and *<u>surgery</u>* without experiencing the cortical contusion (i.e. *uninjured & untreated*). A small number of rats (n=3) in a separate “Naïve” group were *<u>not subjected to any surgery</u>*, to see if the surgical procedures had any effects on the biomarker levels.

All of the remaining groups of rats were exposed to a CCI. Negative control group rats were *injured & untreated*. There were two such groups, one receiving no injection at all (TBI Only) and another group receiving the vehicle buffer being used for Fv-HSP72 (PBS with 5% sucrose, filter sterilized). Vehicle buffer was injected IV 15m post-CCI (TBI & Vehicle) to verify the effects of the vehicle buffer alone. Each rat received a volume of buffer through the tail vein equal to the volume injected into the rats receiving drug. The remaining 6 groups of rats were *injured & treated* IV in the rat tail vein with either a **10 or 30mg/kg dose** of Fv-HSP72 15m post-CCI.

#### Biomarker Tissue Extraction

Shifts in neurodegeneration biomarkers have been reported to peak 48-72h post-TBI^16,17^; hence, all rats were sacrificed at 48h. Brain tissues were removed from both hemispheres of each rat, immediately frozen in liquid nitrogen, stored at -80°C until shipment on dry ice to Rubicon for analysis. JHU provided cortical tissue both from the area of the contusion on the ipsilateral hemisphere and a comparable amount of cortex from the contralateral side. At Rubicon, tissues were transferred to 15 mL conical centrifuge tubes containing 2mL of T-PER™ Tissue Protein Extraction Reagent (ThermoFisher Cat 78510), 5mM EDTA and 1X HALT Protease/Phosphatase Inhibitor Cocktail (ThermoFisher Cat.# PI 78438). Sonication of all samples occurred on ice (10 pulses of 10s each) at 40% power using a Sonics VCX500 with a CV334 needle-nose probe.

#### Biomarker Analysis

Extracts were subjected to a Bradford assay (Pierce™ Coomassie Protein Assay Kit; ThermoFisher Cat. 23200) to determine protein concentrations and to normalize the amount of extract loaded into the wells of a plate. The manufacturer’s provided bovine serum albumin was used for a standard curve to estimate general protein amounts in each sample.

Tissue extracts were plated in triplicate in 96-well plates (Immulon 4 HBX, ThermoFisher) and allowed to adsorb for 1h at 25°C, the extract was then dumped and plates stored at -20°C until use. Prior to analysis, plates were thawed, and wells blocked with blocking buffer (BB: 1% Non-Fat Dry Milk, 1% BSA in TBS, 0.02% Sodium Azide, pH 7.5) for 30m at 25°C. Primary antibodies (**Table 1**) were incubated for 1hr at 25°C, followed by three rinses of each well with TBST (Tris Buffered Saline + Tween-20). Wells were probed with secondary antibodies (**Table 1**) conjugated to horseradish peroxidase for 30m at 25°C, rinsed 3x with TBST, then 2x with TBS before adding TMB (KPL SureBlue Reserve, LGC, Milford, MA). The colorimetric substrate’s conversion from blue to yellow was halted by the addition of a 1% HCl solution and the wells read at the 450nm wavelength using a SpectraMax M5 plate reader (Molecular Devices). To ensure consistency of results across all samples, they were incubated with TMB for the exact same amount of time. The length of TMB incubation was determined by first incubating the plates with the control samples (sham, TBI Only, TBI & Vehicle) in order to determine the length of time before the colorimetric solution approached saturation levels of Abs_450_=2.0 or higher. For some biomarkers, TMB color development was slow. No incubations were allowed to go longer than 15m.

**Table 1.** List of primary (1°) and secondary (2°) antibodies used in our biomarker studies.

| Biomarker | 1°/2° | Host / Antibody | Clone | Supplier | Catalog # | Dilution |
| --- | --- | --- | --- | --- | --- | --- |
| **3-NT** | 1° | Mouse Monoclonal | 39B6 | Sigma Millipore | #SAB5200009 | 1µg/mL |
|  | 2° | Goat anti-Mouse Polyclonal | N/A | Jackson ImmunoResearch | 115-035-166 | 1:1000 |
| All other biomarkers were probed with rabbit primary antibodies and a goat anti-rabbit secondary. |  |  |  |  |  |  |
| **NF-L** | 1° | Rabbit Monoclonal | C28E10 | Cell Signaling | #2837S | 1:2000 |
| **pNF-H** | 1° | Rabbit Polyclonal | N/A | Encor Biotech | RPCA-NF-H | 1:2000 |
| **pTau T181** | 1° | Rabbit Polyclonal | N/A | Abcam | #ab75679 | 1:2000 |
| **pTau T231** | 1° | Rabbit Monoclonal | EPR2488 | Abcam | #ab151559 | 1:5000 |
| **pTau S396** | 1° | Rabbit Monoclonal | EPR2731 | Abcam | #ab109390 | 1:2000 |
| **IL-6** | 1° | Rabbit Monoclonal | C7H24L3 | Invitrogen | #700480 | 1:666 |
|  | 2° | Goat anti-Rabbit Polyclonal | N/A | Jackson ImmunoResearch | 111-036-144 | 1:1000 |

Determination of background absorbances involved plating a single well of each sample extract into an ELISA plate (50µg per sample). All samples were tested by adsorbing for 1h prior to the addition of BB. Wells were then blocked in 200µL of BB for at least 1h and then probed with secondary antibody (**Table 1**) conjugated to horseradish peroxidase for 30m at 25°C, rinsed 3x with TBST, then 2x with TBS before adding TMB. Colorimetric development lasted from 5-15m as reported in the figure legends.

Statistical analysis of biomarker results used an unpaired t-test with Welch’s correction to account for any variances that are unequal between groups. For deviations from unity, a one sample t-test was used. Statistical significance was affirmed with p values ≤ 0.05.

## Results

Our central hypothesis is that the BBB is compromised after a trauma to permit Fv-HSP72 access to regions of tissue damage where it will protect against neurodegeneration and other biological processes that can lead to apoptosis or secondary necrosis. Rubicon Biotechnology has engineered nearly 20 variants of Fv-HSP72 to improve the therapeutic’s efficacy in a microenvironment that is mildly acidic^18,19^ and enriched in reactive radicals^20^. Three of these variants (RBB010, RBB012, RBB012-CTB) were screened in a rat CCI injury model at a 10 or 30mg/kg dose via an IV tail-vein injection 15m post-injury (**Figure 1A**). These dosages were chosen based on a tolerability study that found a no adverse effect limit at the maximum achievable dose of 27.5mg/kg given to the rats (data not shown). Upon improvements in the drug’s purification process, higher dosages were achieved and a high dosage was set at 30mg/kg, with a second dosage at 1/3 the amount (10mg/kg). Fusion of the 3E10 subunit to the HSP72 has been tested with one of two cleavable amino acid linkers. In the CCI studies, one of the variants was tested side-by-side with a cleavable “Swivel” linker (RBB012) or a proprietary “CTB” peptide sequence that is cleaved by cathepsin B (RBB012-CTB), a protease that is upregulated during tissue injury^21^.

**Figure 1.**
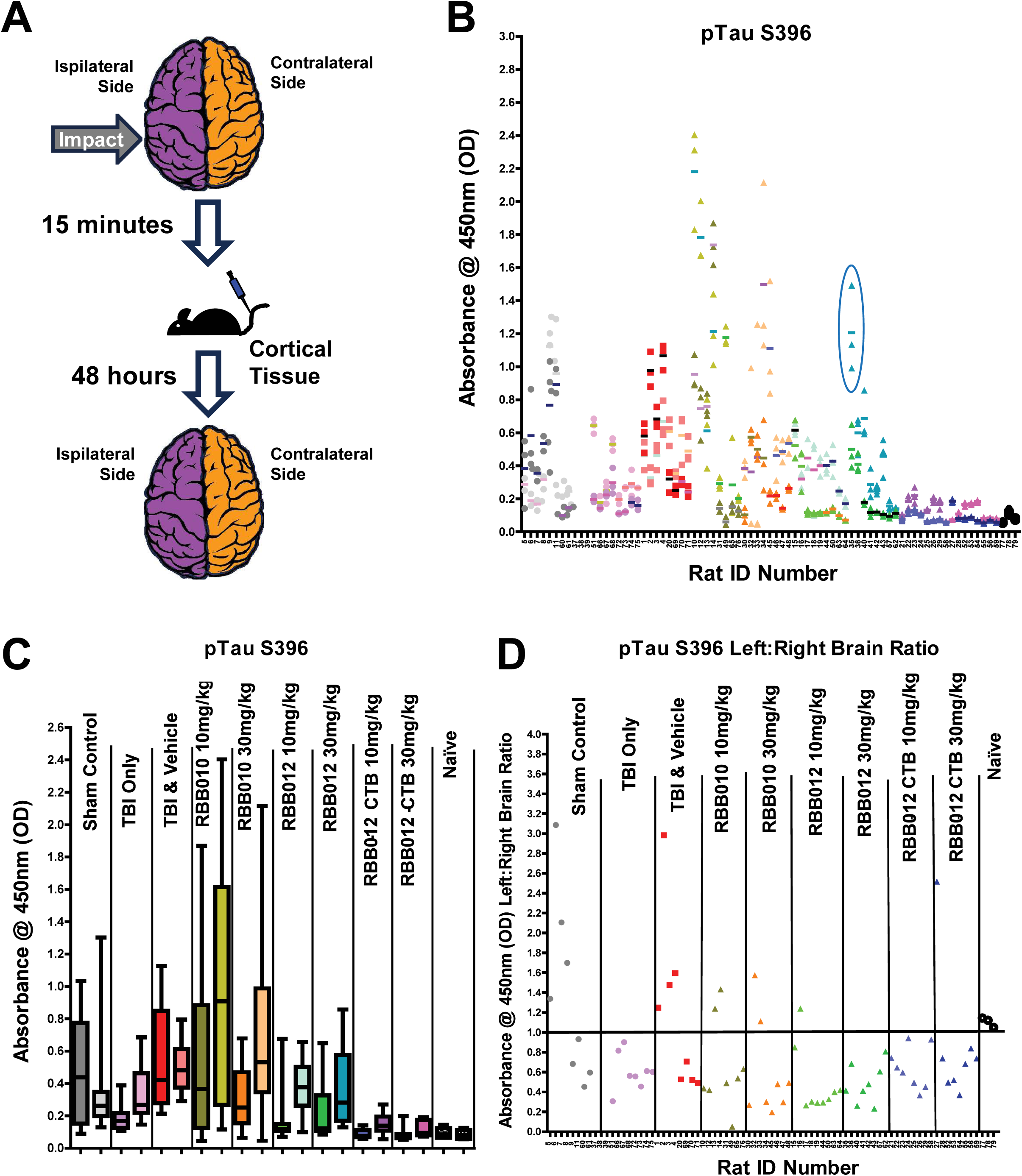
**A)** Schematic of the CCI method. 100µg of cortical tissue extract from each of the rats were plated in triplicate in the wells of an ELISA plate and probed, as described in Methods, at a 1:2000 dilution with a rabbit monoclonal to the pTau S396 phosphorylation site followed by a goat anti-rabbit conjugated to horseradish peroxidase (GAR-HRP). TMB color development was stopped at 5min for this analysis. **B)** A scatter plot showing all of the triplicate results from both the ipsilateral (darker color) and contralateral (lighter color) hemispheres of each rat is a daunting task to read. The mean for the three contralateral absorbance values from Rat #35 were 327.15% greater than the mean for the other contralateral results; hence this data set was dropped from our analysis (circled in blue). **C)** To help visualize the data easier, the same results from each group of rats are presented as box plots with the ipsilateral results on the left and the contralateral on the right. Note that the graphs in (B) and (C) reveal global differences both between treatment groups and within groups (i.e. when comparing ipsi vs. contra). **D)** It is equally important to analyze biomarker differences between the ipsi- and contralateral hemispheres in each rat individually. This is accomplished by plotting the absorbance ratios against a black line denoting an equal distribution of signal in both hemispheres (ipsi / contra = 1). Analysis of the intra-hemispheric differences for each individual rat reveals a lower pTau S396 neurodegeneration signal level in the ipsilateral for rats treated with Fv-HSP72.

Male rats were subjected to a cortical contusion simulating a moderate to severe brain injury. The left hemisphere of the brain was the side that received the contusion; hence, it is the *ipsilateral hemisphere* and the right is the *contralateral* in discussions of the data (**Figure 1A**). Samples were extracted from both hemispheres of the brain 48h post-injury and extracted as described above prior to plating at the same protein quantity, in triplicate. Multiple plates were required to include all of the samples for probing with antibodies to a single biomarker. *To ensure consistency in absorbance values, both ipsi and contra samples from the same rat were always placed next to each other on the same plate.* Wells were developed with colorimetric substrate for the exact same amount of time across multiple plates to accurately compare samples.

### Neurodegeneration Biomarkers

#### Phosphorylated Tubulin Associated Unit (pTau) – T181, T231 and S396

We looked at multiple neurodegeneration markers, including three related to pTau. While phosphorylation at Serine-396 is not a TBI-specific marker, it is strongly implicated in various tau pathologies (particularly Alzheimer’s and to a lesser extent in CTE)^22^. Here, it illustrates the extent biomarker analyses were conducted to screen the 3 Fv-HSP72 variants (**Figure 1B**). Since CCI is a focal injury model, it is incumbent to analyze the contralateral hemisphere to account for the variability among individual rats. Our analysis looked at two phenomena: **1)** Is there a difference in ***INTER***-group biomarker levels between the ipsi- and contra-hemisphere samples *between each group of rats?* For example, are the ipsi results for one group significantly different from the ipsi results in another? **Figure 1C** graphically illustrates the answer. For each group, the box plot on the left corresponds to the ipsi hemisphere and the box on the right corresponds to the contra. **2)** Is there a difference in ***INTRA***-hemispheric biomarker levels between the ipsi- and contralateral samples by analyzing the results *from each individual rat?* **Figure 1D** illustrates the answer by plotting the signal ratio of the ipsi and contra hemispheres (“Left : Right Brain Ratio”). The line at a ratio = 1 represents the condition where the signal is equal in both the left and right hemispheres; hence, suggesting the effects are similar on both sides of the brain. Ratios below this line (i.e. <1) indicate decrease of the biomarker in the ipsilateral hemisphere.

This example is representative of results seen in our other biomarker screenings; overall, RBB010 did not significantly reduce the elevated signals seen in control rats receiving a TBI followed by vehicle buffer (TBI & Vehicle). While RBB010 was screened with most of the remaining biomarkers discussed, it was dropped from further consideration for clinical development. The *inter*-biomarker box plots of the ipsilateral data groups clearly showed a statistically significant drop in the neurodegenerative marker pTau S396 (*p<0.0001*) for rats treated either with RBB012 or RBB012-CTB, regardless of the 10 or 30mg/kg dose, when compared to their ipsi-counterpart in the TBI & Vehicle group (**Figure 1C**). Similarly, the *inter*-biomarker box plot comparison of the contralateral groups showed significant drops in absorbance values compared to the TBI & Vehicle group (RBB012: 10mg/kg *p=0.0053* and 30mg/kg *p=0.0291*; RBB012-CTB: 10 and 30mg/kg *p<0.0001*).

Drug localization to the injury site can result in differences in the drug’s effect on the ipsi-versus contralateral hemispheres (in **Figure 1C** compare the left box plot to its right counterpart in each group). For pTau S396, this effect is seen with the significant decreases of biomarker signals in the ipsilateral vs contralateral within each group (RBB012: 10mg/kg *p<0.0001* and 30mg/kg *p=0.0075*; RBB012-CTB: 10mg/kg *p<0.0001* and 30mg/kg *p=0.0127)*. This difference was not significant in the TBI + Vehicle group (*p=0.4172*). To further characterize these *intra*-biomarker differences, an analysis of *each individual rat* reveals a drop in biomarker levels for rats treated with Fv-HSP72 below unity (ipsi/contra =1; **Figure 1D**). Deviations from unity can be statistically analyzed with a two-tailed t-test and reveal significance in 3 of the four Fv-HSP72 treatments (RBB012: 10mg/kg *p=0.0017* and 30mg/kg *p=0.0008*; RBB012-CTB: 10mg/kg *p=0.0022*). The importance of looking at both inter-group and intra-hemispheric differences is highlighted in the fact that treatment with RBB012-CTB at 30mg/kg did not significantly deviate from unity (p=0.5878); yet, **Figure 1C** clearly shows the effect of the drug at that dosage was to lower biomarker signals in both hemispheres down to levels comparable to naïve rats that were not even surgically manipulated.

Unlike RBB010 and RBB012 (both possessing a Swivel linker), RBB012-CTB proved to be the most consistently efficacious with respect to our biomarker screens. Possessing a cathepsin B cleavable linker, the results persuaded us that RBB012-CTB had the best chance of success when tested in clinical trials. We focus our remaining presentation on this variant.

Tau phosphorylation at residues **Threonine-231 (T231)**^23^ and **Threonine-181 (T181)**^24^ have become important biomarkers for the early prognosis of traumatic brain injury severity. For pTau T231, both dosages of RBB012-CTB reduced the neurodegenerative marker significantly in the ipsilateral hemisphere, but not the contralateral, when compared to the same hemispheres in those rats given vehicle buffer post-TBI (**Figures 2A,B**). This decrease in the neurodegeneration signal for the ipsi-hemisphere was statistically significant with the inter- or intra-biomarker analysis (**Figures 2B and 2C**, respectively). For pTau T181, the two dosages had to be analyzed against the TBI+Vehicle control on separate dates, but the trends were similar to T231 (**Figure 3**). At both dosages, inter-group differences between the ipsi-samples are significant compared to the TBI + Vehicle control (**Figures 3C,D**). Intra-hemispheric differences were also significant (**Figures 3C-F**).

**Figure 2.**
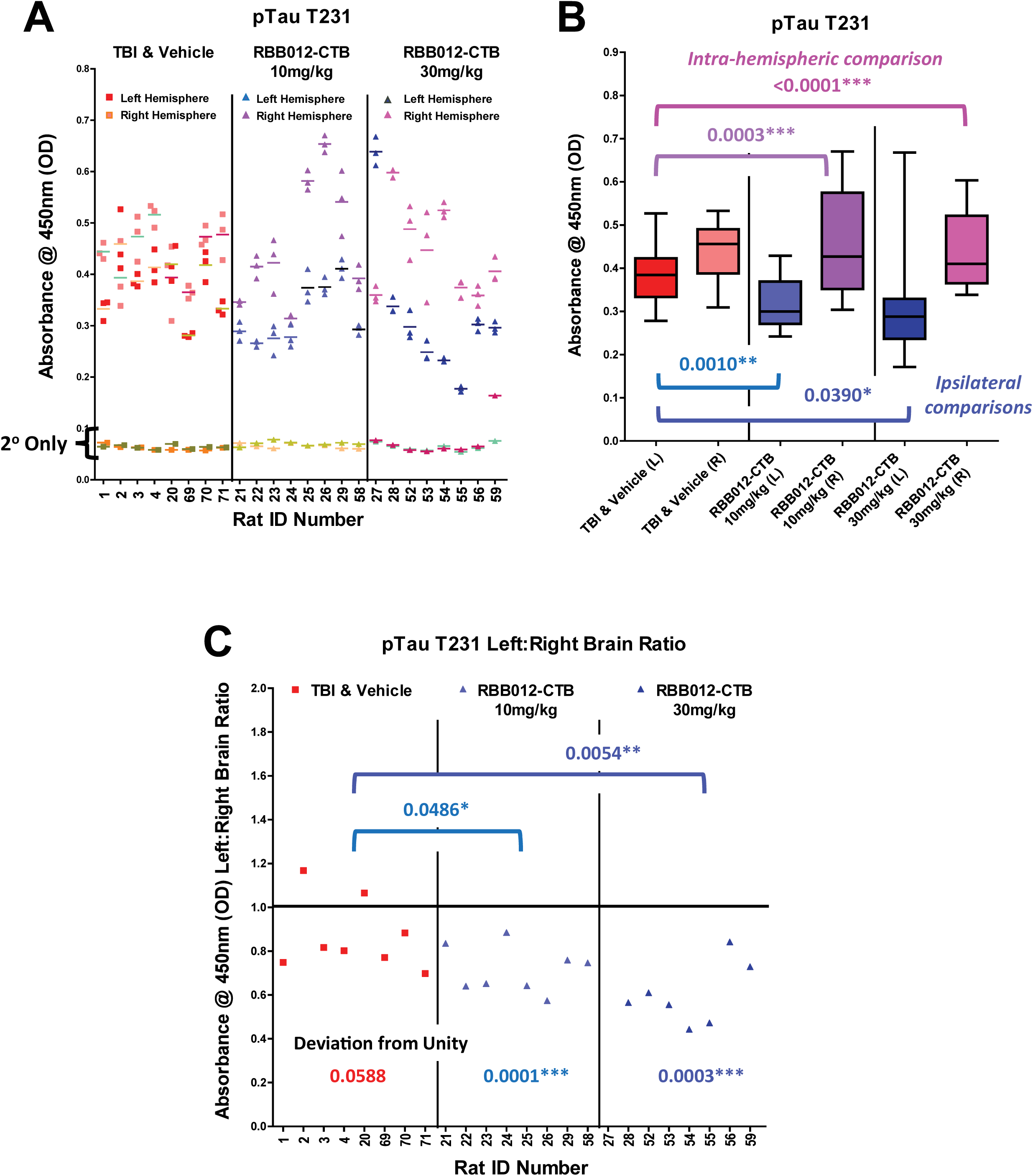
**A)** 100µg of cortical tissue extract from each of the rats were plated in triplicate in the wells of an ELISA plate and probed, as described in Methods, at a 1:5000 dilution with a rabbit monoclonal to the pTau T231 phosphorylation site followed by a GAR-HRP secondary. Baseline absorbances (2° Only) were obtained in wells probed only with GAR-HRP as described in Methods. TMB color development was stopped at 7.5m for this analysis. All wells probed with primary antibody resulted in signals well above background. **B)** The same data set presented as box plots show statistically significant inter-group decreases in the ipsilateral hemispheres of those rats treated with RBB012-CTB versus buffer only post-CCI (10mg/kg p=0.0010; 30mg/kg p=0.0390). In fact, the intra-hemispheric absorbances from the ipsi-samples were significantly lower than the contra within both treatment groups (10mg/kg p<0.0001; 30mg/kg p=0.0003). **C)** This intra-hemispheric difference can also be analyzed as ratios for each rat in order to normalize any variation. Again, the rats receiving drug post-CCI deviated from unity (10mg/kg p=0.0001; 30mg/kg p=0.0003) and from the vehicle control (10mg/kg p=0.0486; 30mg/kg p=0.0054) significantly.

**Figure 3.**
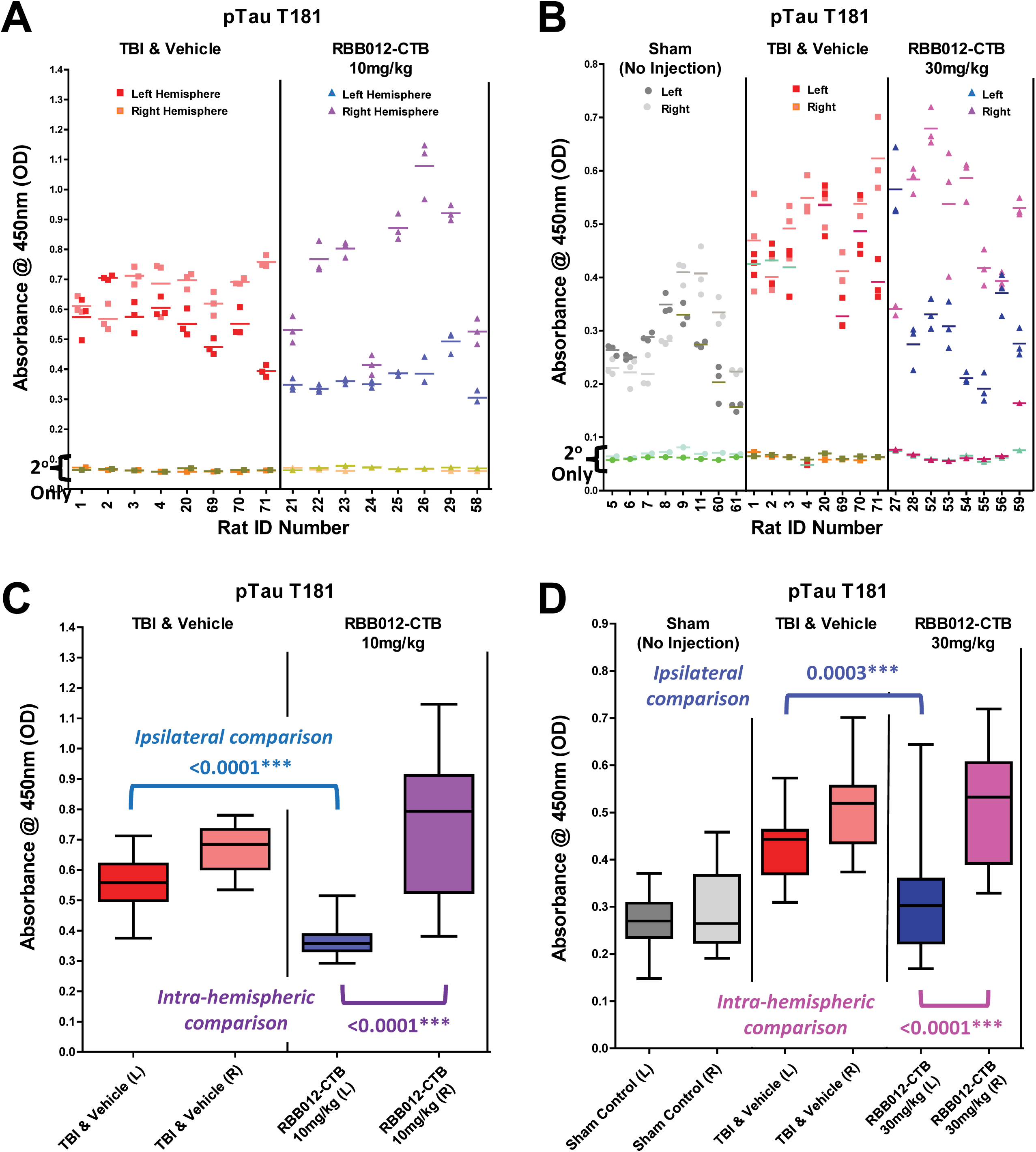
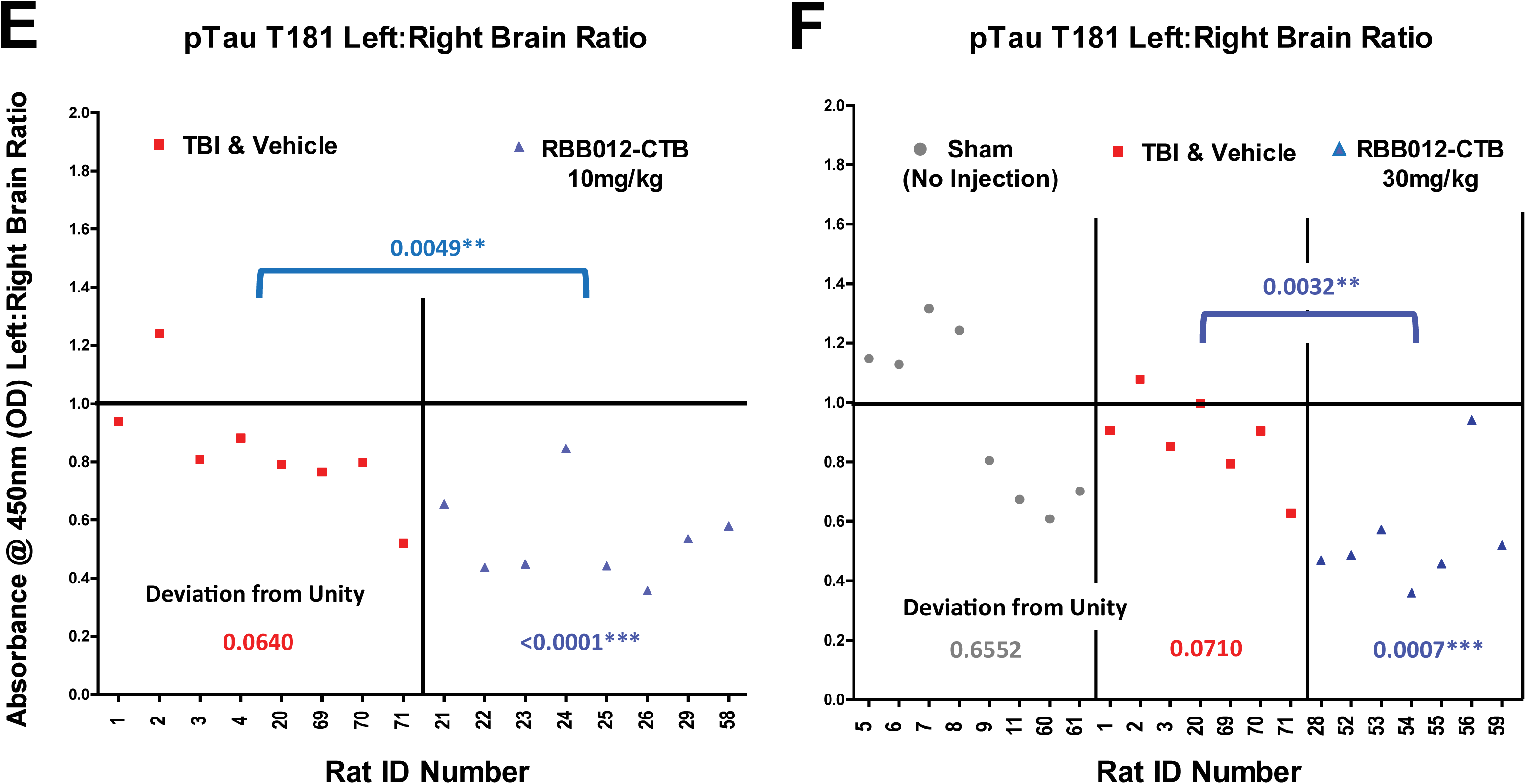
**A)** 100µg of cortical tissue extract from each of the rats were plated in triplicate and probed as described in Methods, at a 1:2000 dilution with a rabbit polyclonal to the pTau T181 site followed by a GAR-HRP secondary. TMB color development for the **A)** 10mg/kg sample analysis was stopped at 5m and for **B)** 30mg/kg at 12.5m. Baseline absorbances (2° Only) were obtained in wells probed only with GAR-HRP as described in Methods and color developed for 7.5m. The same data sets presented as box plots show highly significant inter-group and intra-hemispheric decreases in the ipsilateral samples of those rats treated with RBB012-CTB versus vehicle only post-CCI. **C)** 10mg/kg both, inter-group and intra-hemispheric: p<0.0001. **D)** 30mg/kg inter-group: p=0.0003; intra-hemispheric: p<0.0001). Breaking the intra-hemispheric analysis down further to account for individual rat variability still shows significant deviation from unity and the vehicle controls for **E)** 10mg/kg: p<0.0001, p=0.0049, respectively and **F)** 30mg/kg: p=0.0007, p= 0.0032, respectively.

#### Neurofilament biomarkers (NF-L, pNFH)

Neurofilaments (NF) are cytoskeletal structural proteins released as a result of axonal damage. The NF-Light Chain protein fragment (NF-L) is becoming an important diagnostic for determining neurodegeneration in blood samples^24^. The phosphorylated form of the NF-Heavy Chain fragment (pNFH) is also considered a useful marker. Due to the focal nature of the CCI injury model, our biomarker analysis was confined to neuronal tissue. Both dosages of RBB012-CTB decreased the NF-L content in ipsi-samples compared to those rats receiving only a vehicle control post-CCI (**Figures 4C,D**). In fact, NF-L levels in Fv-HSP72 rats approached those seen in Sham rats exposed only to the surgical procedure (**Figure 4D**) and Naïve rats that were not even surgically manipulated (**Figure 4C**). Individually, most rats treated with either dose of RBB012-CTB had lower NF degeneration in the form of NF-L peptide in the ipsilateral versus contralateral hemisphere (**Figures 4E,F**). In the case of pNF-H, the 10mg/kg dose was similar to the vehicle control (**Figure 5C**) and a significant difference was only seen at 30mg/kg (**Figure 5D**). The 30mg/kg dose results reaffirm the importance of comparing intra-hemispheric differences at both the group and individual levels; pNF-H levels were decreased significantly in both hemispheres for rats receiving the drug (**Fig. 5D** p<0.0001) to such an extent that individual rats do not show significant deviations from unity (**Fig. 5F** p=0.2630).

**Figure 4.**
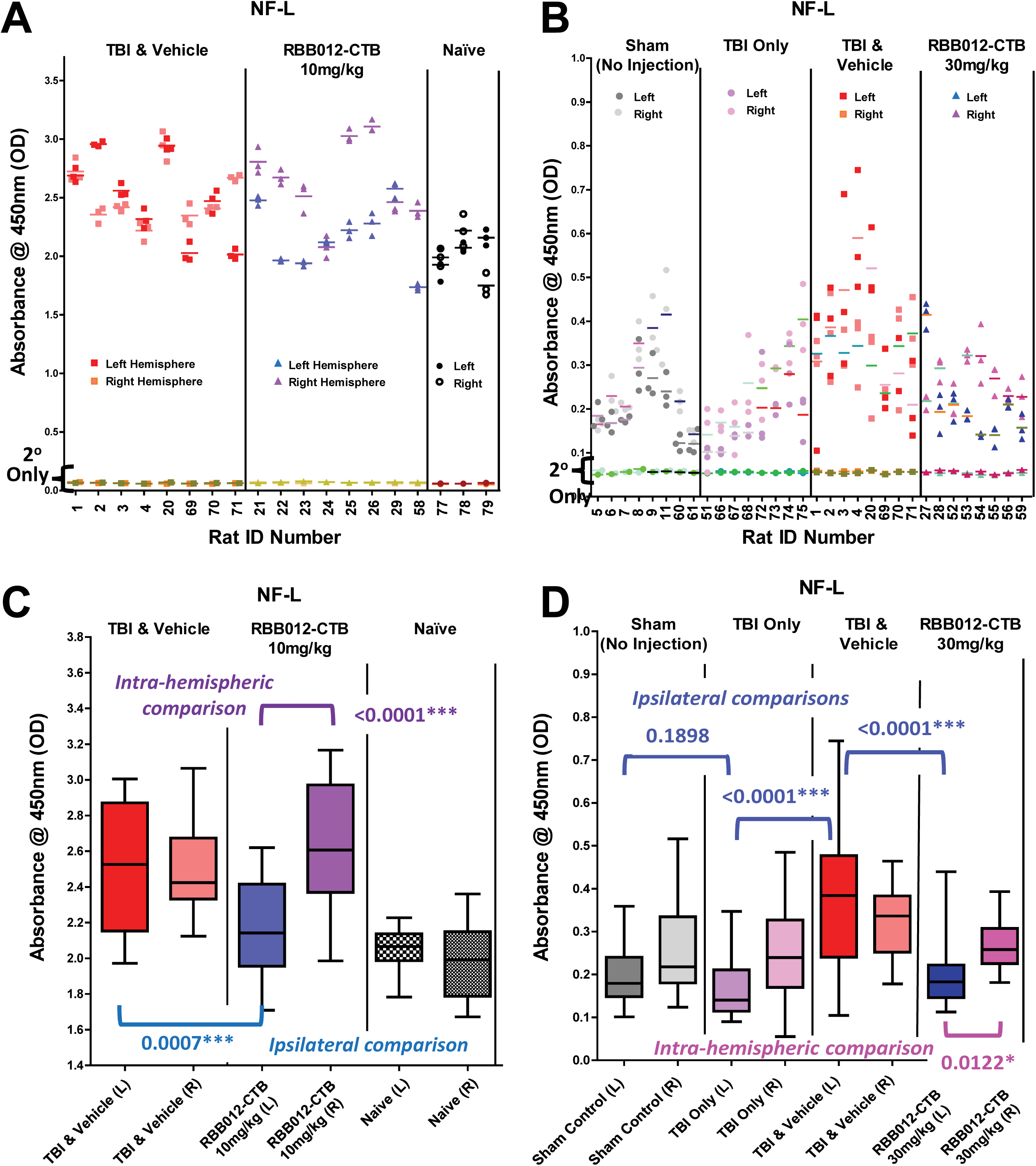
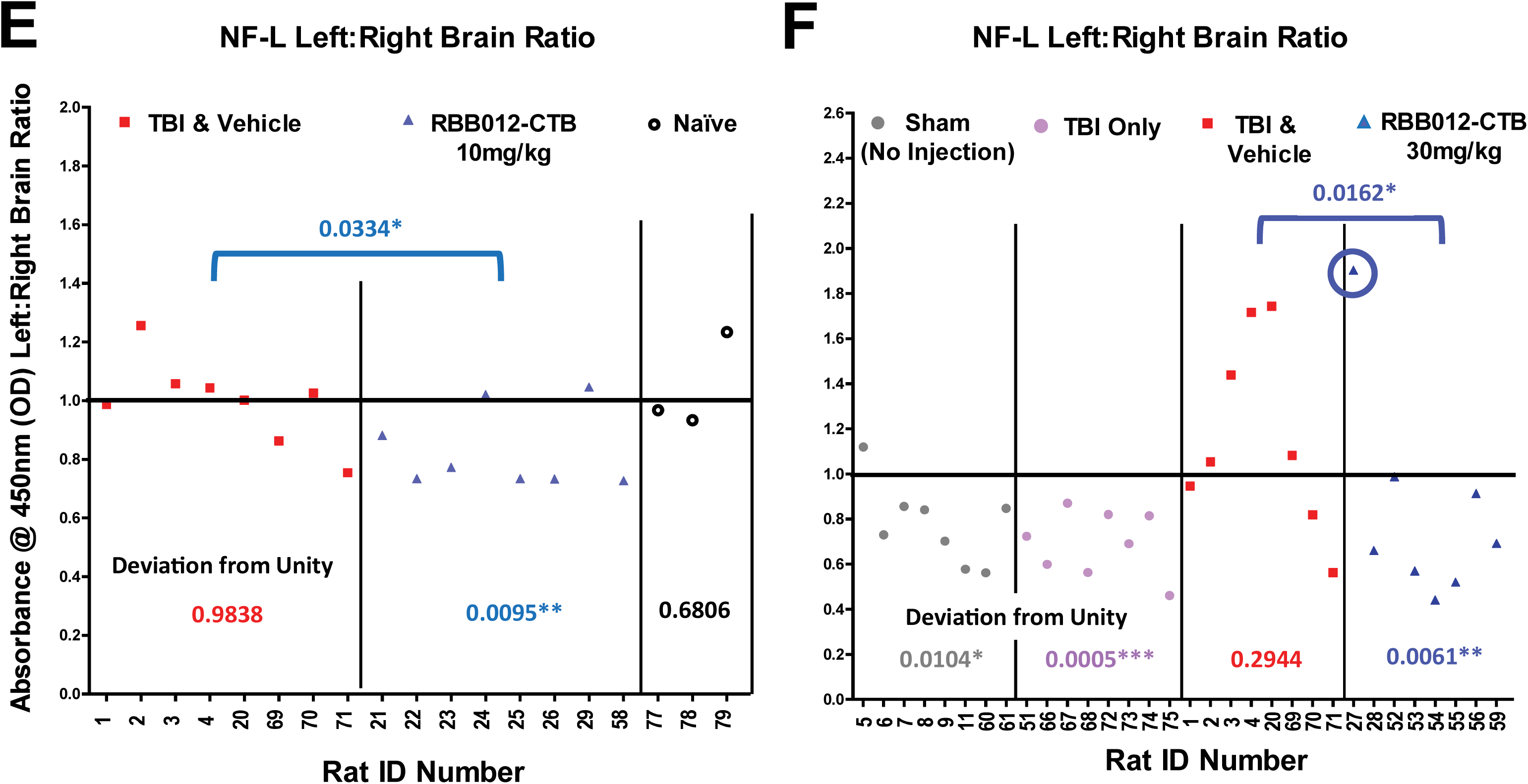
**A)** 100µg of cortical tissue extract from each of the rats were plated in triplicate and probed as described in Methods, at a 1:2000 dilution with a rabbit monoclonal to NF-L followed by a GAR-HRP secondary. TMB color development for the **A)** 10mg/kg and **B)** 30mg/kg samples analysis was stopped at 10m. Baseline absorbances (2° Only) were obtained in wells probed only with GAR-HRP as described in Methods and color developed for **A)** 7.5m and **B)** 10m. The same data sets presented as box plots show highly significant inter-group and intra-hemispheric decreases in the ipsilateral samples of those rats treated with RBB012-CTB versus vehicle only post-CCI. **C)** 10mg/kg inter-group: p=0.0007 and intra-hemispheric: p<0.0001. **D)** 30mg/kg inter-group: p<0.0001; intra-hemispheric: p=0.0122. Breaking the intra-hemispheric analysis down further to account for individual rat variability still shows significant deviation from unity and the vehicle controls for **E)** 10mg/kg: p=0.0334, p=0.0095, respectively and, in the case of **F)** 30mg/kg: p=0.0061, p= 0.0162, when the 278.33% outlier for Rat #27 is dropped (circled in blue).

**Figure 5.**
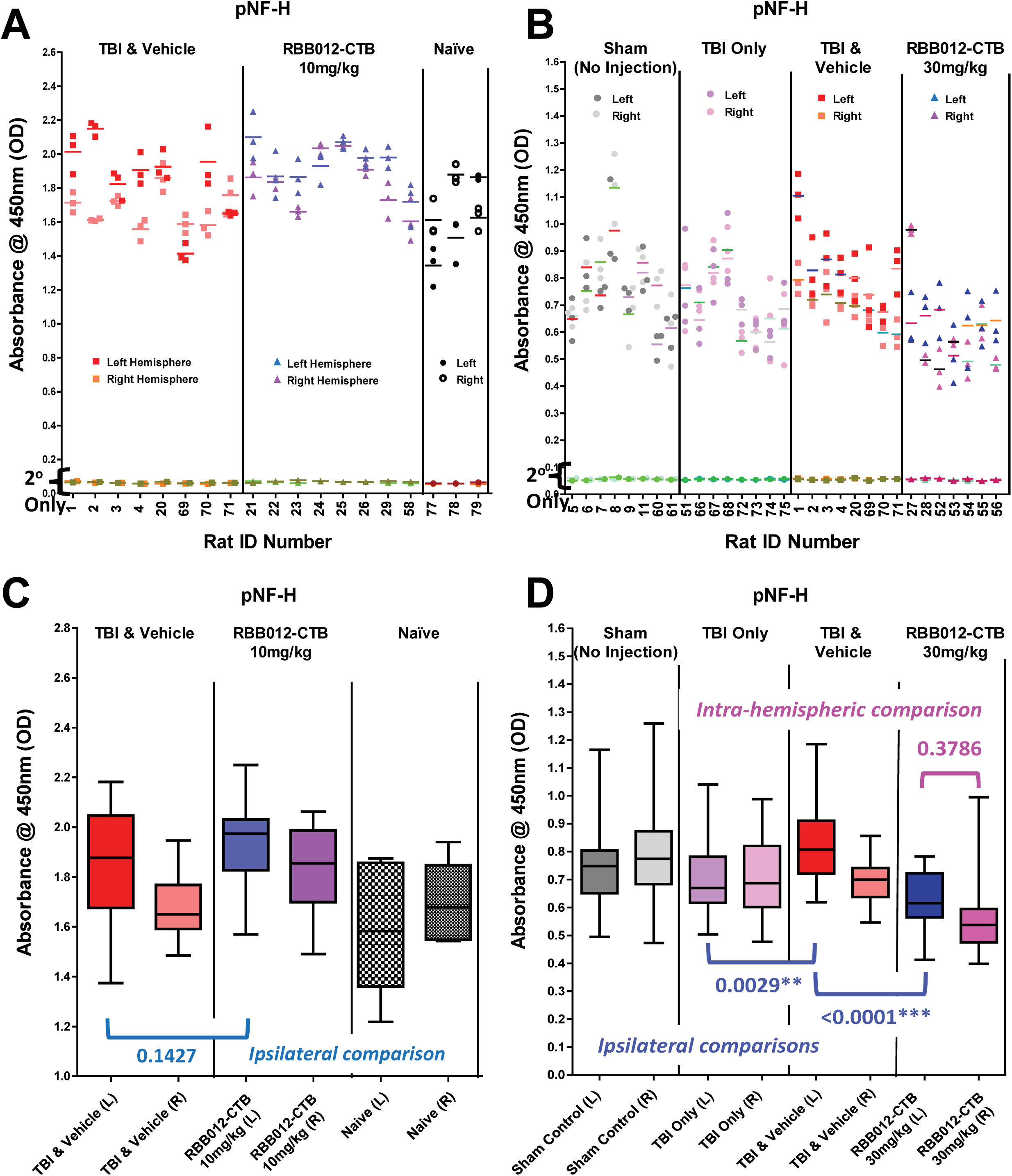
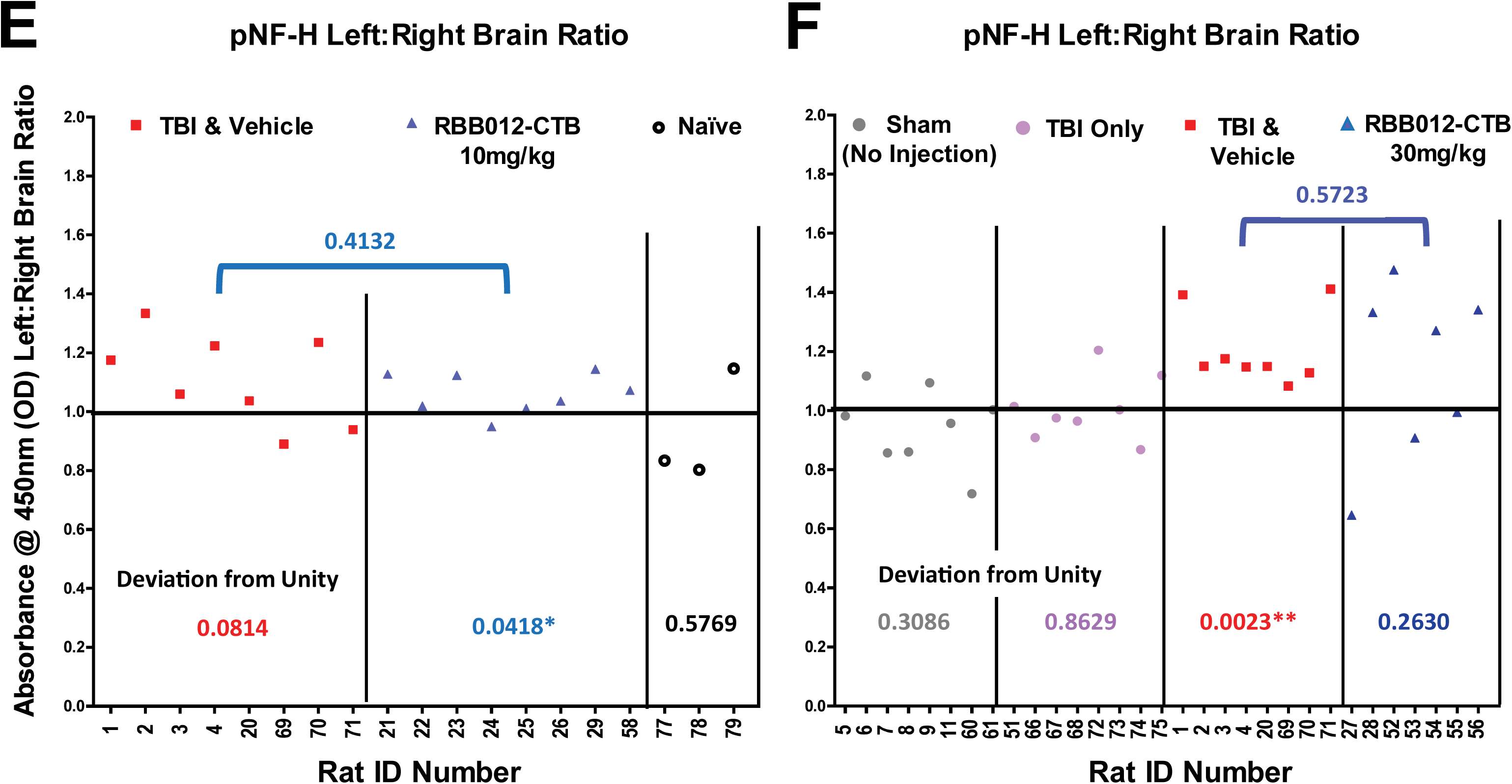
**A)** 100µg of cortical tissue extract from each of the rats were plated in triplicate and probed as described in Methods, at a 1:2000 dilution with a rabbit polyclonal to the pNF-H site followed by a GAR-HRP secondary. TMB color development for the **A)** 10mg/kg and **B)** 30mg/kg samples analysis was stopped at 2m. Baseline absorbances (2° Only) were obtained in wells probed only with GAR-HRP as described in Methods and color developed for **A)** 7.5m and **B)** 10m. **C)** The same data sets presented as box plots show no improvement over the vehicle control for this biomarker at 10mg/kg. **D)** However, significant decreases occur in both the ipsi- and contra-hemispheres at 30mg/kg inter-group: p<0.0001. **E)** Breaking the intra-hemispheric analysis down further to account for individual rat variability shows a deviation from unity for the 10mg/kg dose (p=0.0418), but that is actually due to a slight increase in the pNF-H absorbance of the ipsilateral samples, and this difference is not significant when compared to the vehicle controls (p=0.4132). **F)** Decreases of pNF-H in both hemispheres for the rats dosed at 30mg/kg are not reflected as deviations from unity for the left:right hemisphere brain ratio analysis.

Note, in some of the biomarker studies, absorbance values were greater in rats receiving Vehicle buffer versus rats receiving a TBI Only with no buffer injection (**Fig 4D** p<0.0001, **Fig 5D** p=0.0029); in fact, TBI Only rats can be statistically similar to Sham animals (**Fig 4D** p=0.1898). One possible reason for this could be buffer injections cause reperfusion of fluid flow into the crushed tissues resulting in increased oxidative stress and tissue damage, not unlike the reperfusion injury seen after removal of a vascular occlusion. This hypothesis is supported by observations in parallel studies of blast injured rats, which did not receive a direct impact, and had similar absorbance values between TBI Only and TBI+Vehicle animals (manuscript in preparation). Thus, in these CCI studies, the most appropriate control to compare against our Fv-HSP72 treated animals are the rats receiving Vehicle buffer without drug.

### Oxidative Stress

The secondary processes following a TBI include excessive intracellular calcium (Ca^+2^) accumulation leading to mitochondrial dysfunction, excitotoxicity and production of reactive oxygen and nitrogen species (ROS/ RNS). Several biomarkers are known for measuring levels of tissue damage caused by these free radicals; in our case, measuring the nitration of tyrosine residues as it accumulated on proteins as 3-nitrotyrosine (**3-NT**) proved the most useful for analysis of cortical tissue (**Figure 6**). Nitration occurs when the hydroxyl group of the tyrosine reacts with RNS, such as peroxynitrite formed from the reaction of nitric oxide and superoxide. Nitration levels dropped dramatically to the same level as naïve rats and both dosages were sufficient to affect tissues in the two hemispheres of the brain (**Figures 6A,B**). At 30mg/kg, the decrease in the ipsilateral was even greater when accounting for the variability in each individual rat (**Figure 6C**, p=0.0308).

**Figure 6.**
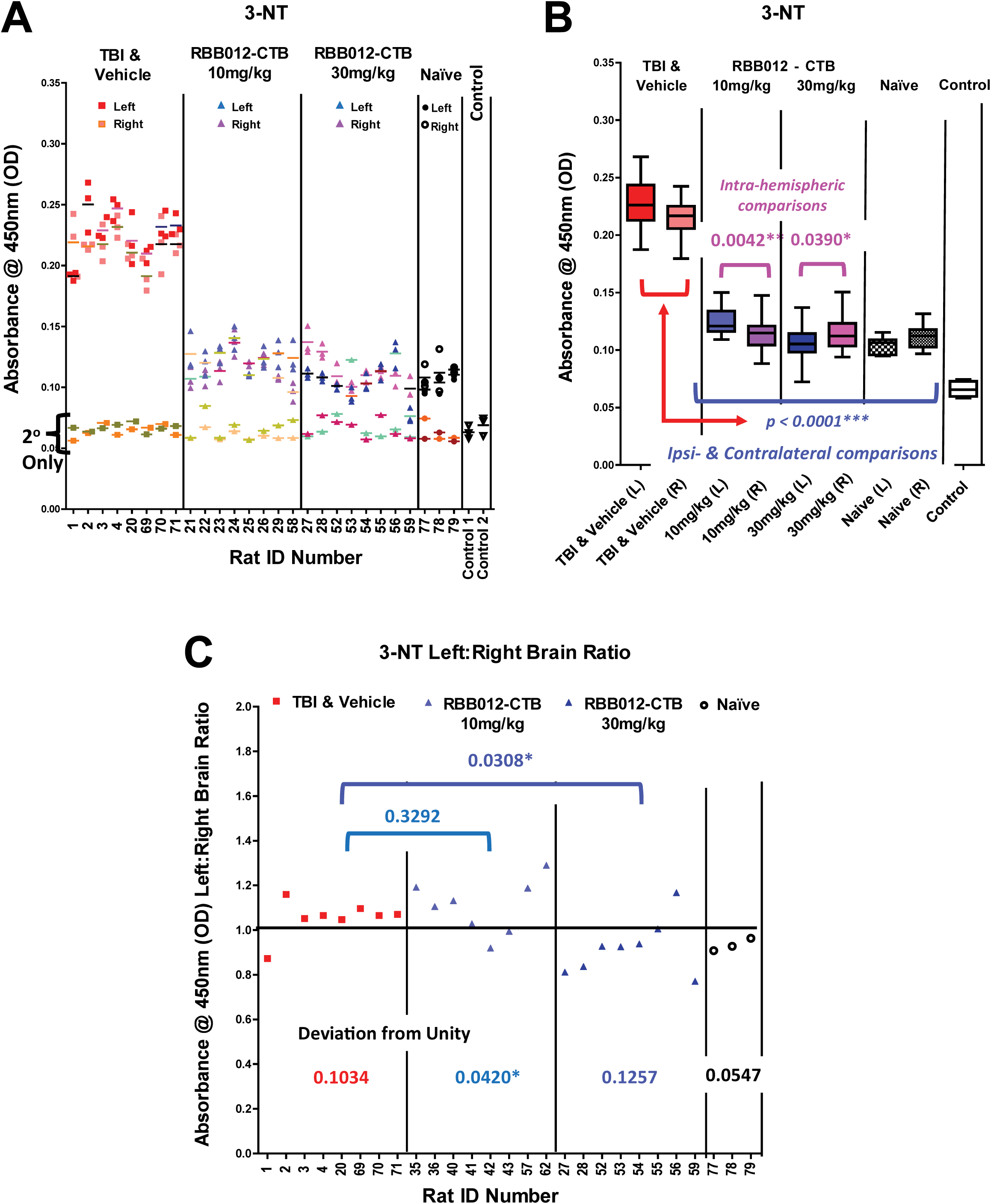
**A)** 100µg of cortical tissue extract from each of the rats were plated in triplicate in the wells of an ELISA plate and probed, as described in Methods, at a 1µg/mL dilution with a mouse monoclonal to the 3-NT adduct followed by a goat anti-mouse conjugated to horseradish peroxidase (GAM-HRP). TMB color development was stopped at 15m for all wells probed with primary antibody, including 6 “control” wells (3 wells in two separate rows) that had no extract at all. Baseline absorbances (2° Only) were obtained in wells probed only with GAM-HRP as described in Methods, with color development stopped at 10m once it was clear the 2° Only signal was not going to rise above the absorbances obtained for the control wells. **B)** The same data set presented as box plots show statistically significant inter-group decreases in both the ipsilateral and contralateral hemispheres of those rats treated with RBB012-CTB, regardless of dosage, versus Vehicle buffer only (p<0.0001). Absorbances were similar to those obtained from naïve rats that were not surgically manipulated at all. The intra-hemispheric absorbances from the ipsi-samples were significantly different from the contra within both treatment groups (10mg/kg p=0.0042; 30mg/kg p=0.0390). **C)** This intra-hemispheric difference can also be analyzed as ratios for each rat in order to normalize any variation. A deviation from unity for the 10mg/kg dose (p=0.0420) is actually due to a slight increase in the 3-NT absorbance of the ipsilateral samples, but this difference is not significant when compared to the vehicle controls (p=0.3292). Decreases of 3-NT in both hemispheres for the rats dosed at 30mg/kg are not reflected as deviations from unity for the left:right hemisphere brain ratio analysis (p=0.1257). Although the absorbances seen in both hemispheres are far lower than that seen in the TBI & Vehicle group, the ipsilateral hemispheres at 30mg/kg are still significantly lower than their contralateral counterparts when comparing to the Vehicle controls (p=0.0308).

### Neuroinflammation

Although not specific to TBI injuries, proinflammatory cytokines, such as IL-6, are released into brain tissue and blood presumably by reactive glial cells post-injury^16^. IL-6 is considered a subacute biomarker, increasing in the blood for days and weeks post-impact^16^. Since tissues were harvested 48 hours post-injury, it made for an appropriate marker to investigate (**Figure 7**). Unlike 3-NT, the 10mg/kg dose did not reduce IL-6 levels in the ipsilateral (**Figure 7C**). Intra-hemispheric differences were statistically significant (**Figures 7C,E**), but this was due to an anomalous increase in IL-6 levels in the contralateral hemisphere of those rats receiving RBB012-CTB. The 30mg/kg dose dropped IL-6 levels in both hemispheres to those seen in naïve rats (**Figure 7D**); a comparison of the ipsilateral absorbances between the two groups being statistically similar (p=0.1022). Intra-hemispheric differences were also significant with the ipsilateral absorbances lower when compared inter-group (p=0.0001) or individual rat (**Figure 7F**, p=0.0056). This example again illustrates the need to look at group and individual results. While **Figure 7F** suggests no significance in the ipsi:contra brain ratios of TBI & Vehicle versus those rats receiving RBB012-CTB at 30mg/kg (p=0.4554), **Figure 7D** reveals a highly significant effect on the ipsilateral absorbances between the two groups (p<0.0001).

**Figure 7.**
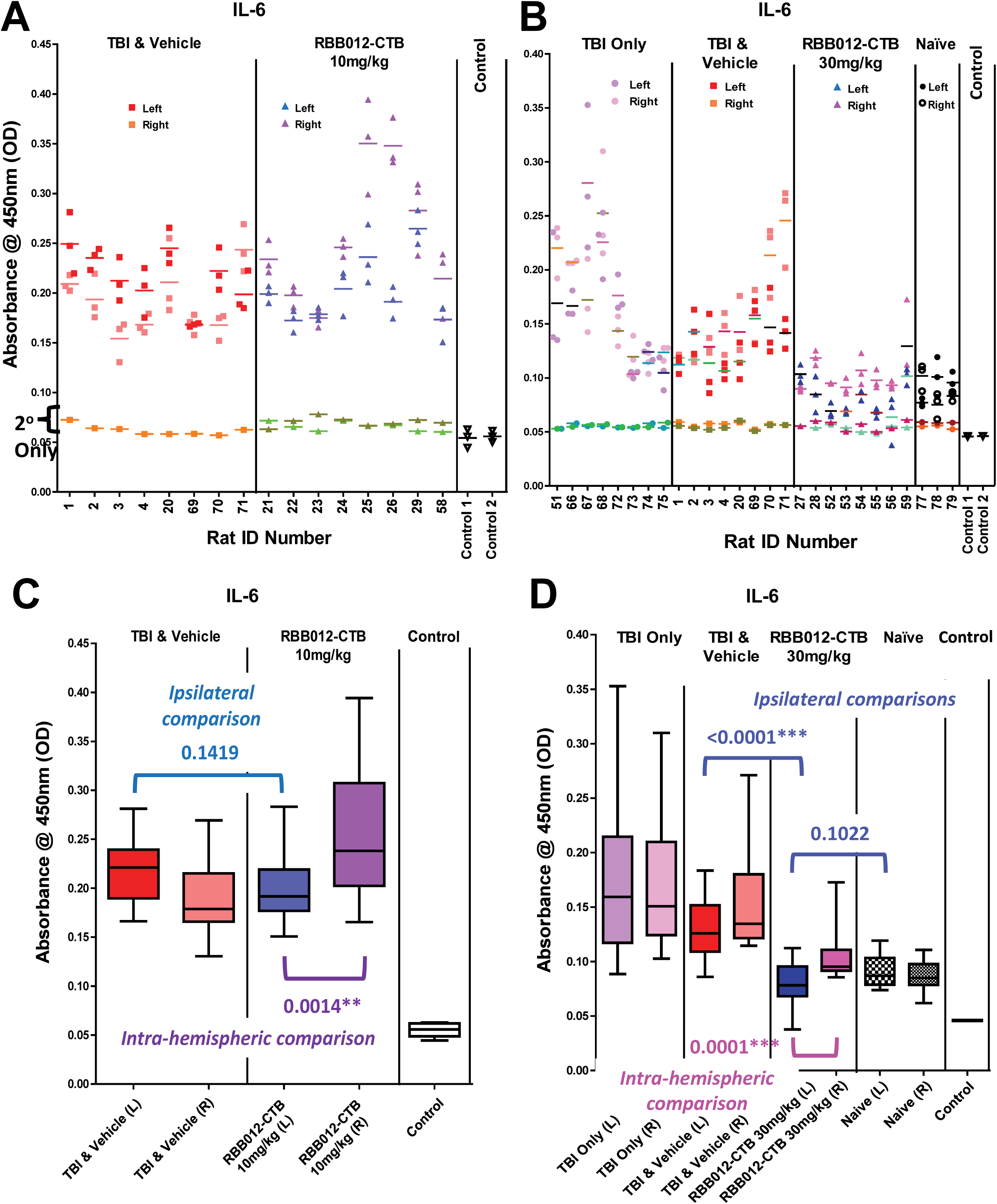
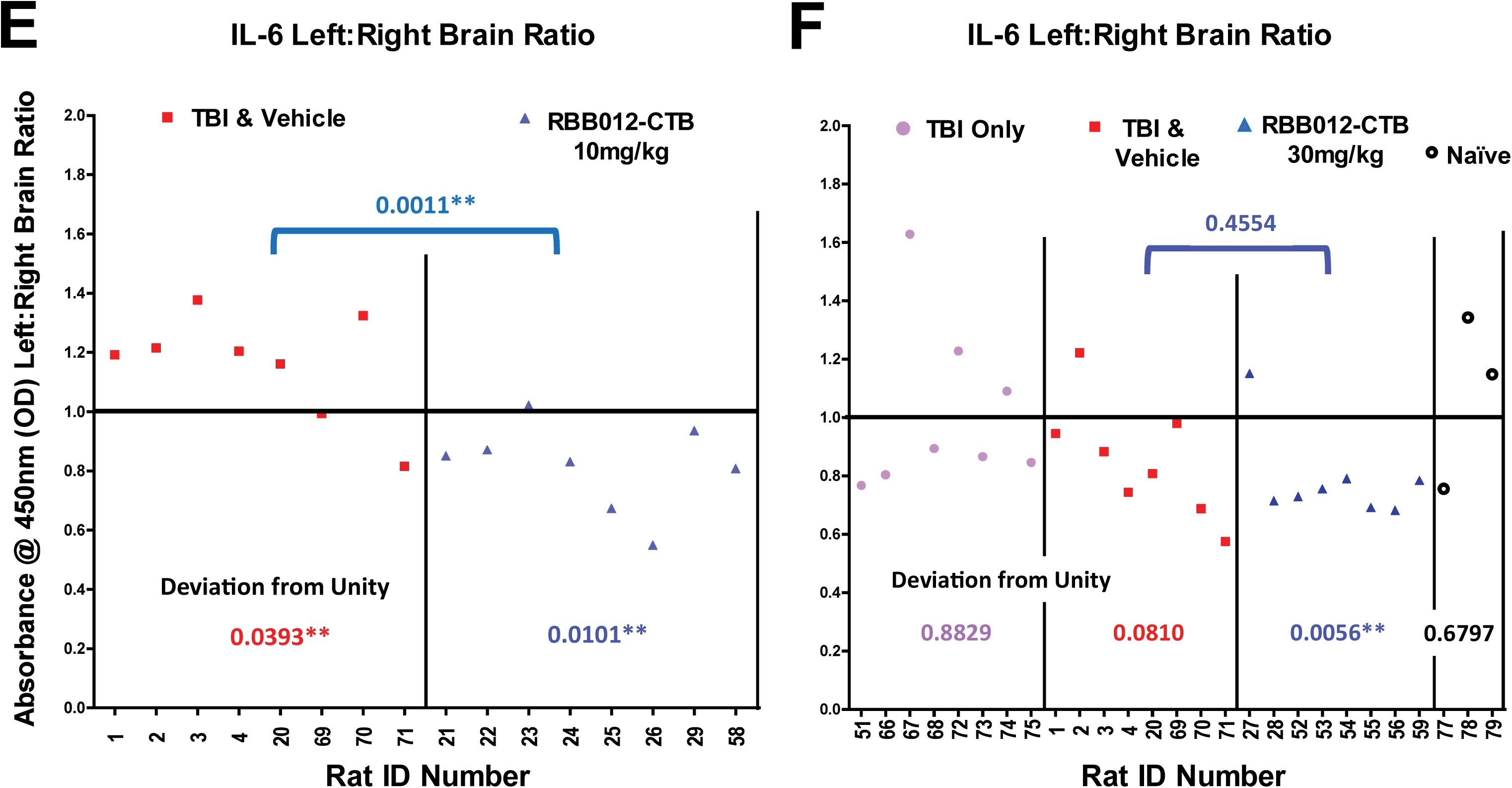
**A)** 100µg of cortical tissue extract from each of the rats were plated in triplicate and probed as described in Methods, at a 1:666 dilution with a rabbit polyclonal to IL-6 followed by a GAR-HRP secondary. TMB color development for the **A)** 10mg/kg and **B)** 30mg/kg samples analysis was stopped at 10m for all wells probed with primary antibody, including 6 “control” wells (3 wells in two separate rows) that had no extract at all. Baseline absorbances (2° Only) were obtained in wells probed only with GAR-HRP as described in Methods, and color developed for **A)** 7.5m and **B)** 10m (once it was clear the 2° Only signal was not going to rise above those of the control wells). **C)** Box plots show no improvement over the vehicle control for this biomarker at 10mg/kg (p=0.1419). **D)** However, significant decreases occur at 30mg/kg inter-group ipsilateral TBI&Vehicle vs RBB012-CTB comparison: p<0.0001. The IL-6 levels drop to those in naïve rats (p=0.1022). **E)** Breaking the intra-hemispheric analysis down further to account for individual rat variability shows a deviation from unity for the 10mg/kg dose (p=0.0101), but that is actually due to an increase in the IL-6 absorbance of the contralateral samples, the increase being significant when compared to the vehicle controls (p=0.0011). **F)** Decreases of IL-6 occurred in both hemispheres for the rats dosed at 30mg/kg with the ipsilateral samples in nearly all of the rats having lower absorbances as reflected in the deviations from unity (p=0.0056).

### Drug Clearance in the Blood

To prove drug uptake into the brain, it was critical to distinguish the modified HSP72 constituting the three Fv-HSP72 variants from endogenous HSP72 induction in the brain. Proprietary primary sequence variations to the HSP72 provided several tryptic peptide sequences that were detectable by MS, of which some are highlighted in a schematic (**Figure 8A**). Selection of these MS-based biomarkers is discussed in Methods.

**Figure 8.**
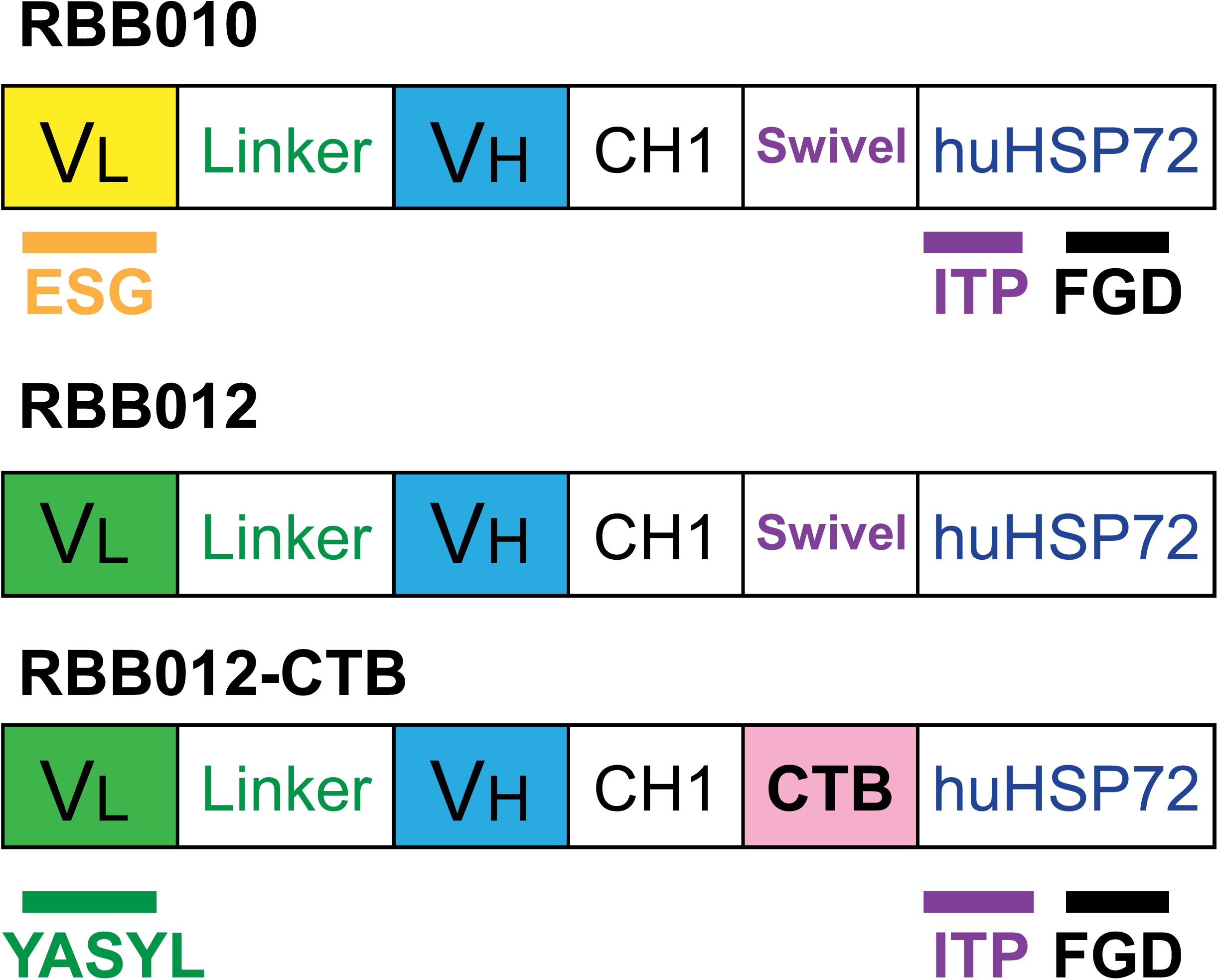
Schematics of the 3 Fv-HSP72 variants tested. The ESG and YASYL peptides allow for detection of the 3E10 scFv subunit consisting of a variable light chain (V_L_), linker, variable heavy chain (V_H_), and a portion of a human constant heavy chain (CH1). These peptides also allow us to differentiate the RBB010 from RBB012 variants by mass spec because of a point mutation in the V_L_ sequence. C-terminal to the 3E10 subunit are the clevable linkers (Swivel, CTB) followed by a modified human HSP72 common to all 3 variants. The ITP and FGD peptides track this modified HSP72 and differentiate it from endogenous HSP72 in tissue extracts. The ITP peptide detects Fv-HSP72 regardless of the oxidative state of the tissues it is in. FGD has a methionine residue that can be oxidized and allows us to evaluate the relative oxidation level of our drug by setting the mass spectrometer to detect FGD peptide in three states of oxidation (methionine unoxidized, methionine sulfoxide and methionine sulfone).

PK studies tracking both the 3E10 and HSP72 subunits in our fusion protein involved IV tail vein injections into rats either 1) exposed to anesthesia, surgery and a CCI or 2) naïve rats only exposed to anesthesia. Both, naïve and TBI-exposed rats were divided into 3 groups (n=5/group) with one group sacrificed 1h post-drug injection, and two other groups at 4h and 12h. All 3 Fv-HSP72s were subjected to PK analysis of the brain and plasma tissues, making for a total of 18 groups. Hundreds of tissue analyses were conducted, but we focus here on the RBB012-CTB results. The plasma clearance of RBB012-CTB injected IV 15m after exposure are plotted both as a composite of 3 peptide biomarkers (**Figure 9A**) and individually to illustrate the Lower Limit of Quantitation (LLOQ), also known as the signal to noise (S/N) threshold, whose determination is discussed below (**Figures 9B-D**). Both injured and naïve rats have a plasma concentration of the YASYL peptide, and presumably the 3E10 subdomain of RBB012-CTB that it represents, below this limit of accurate quantitation 12h after IV injection. Accurate quantitation of the ITP and FGD peptides occurs for the first 4h before dropping below their LLOQs sometime before the 12h mark. Based on our calculation methods, the ITP’s is low (1.95 fmoles/µL) and the FGD’s is essentially zero.

**Figure 9.**
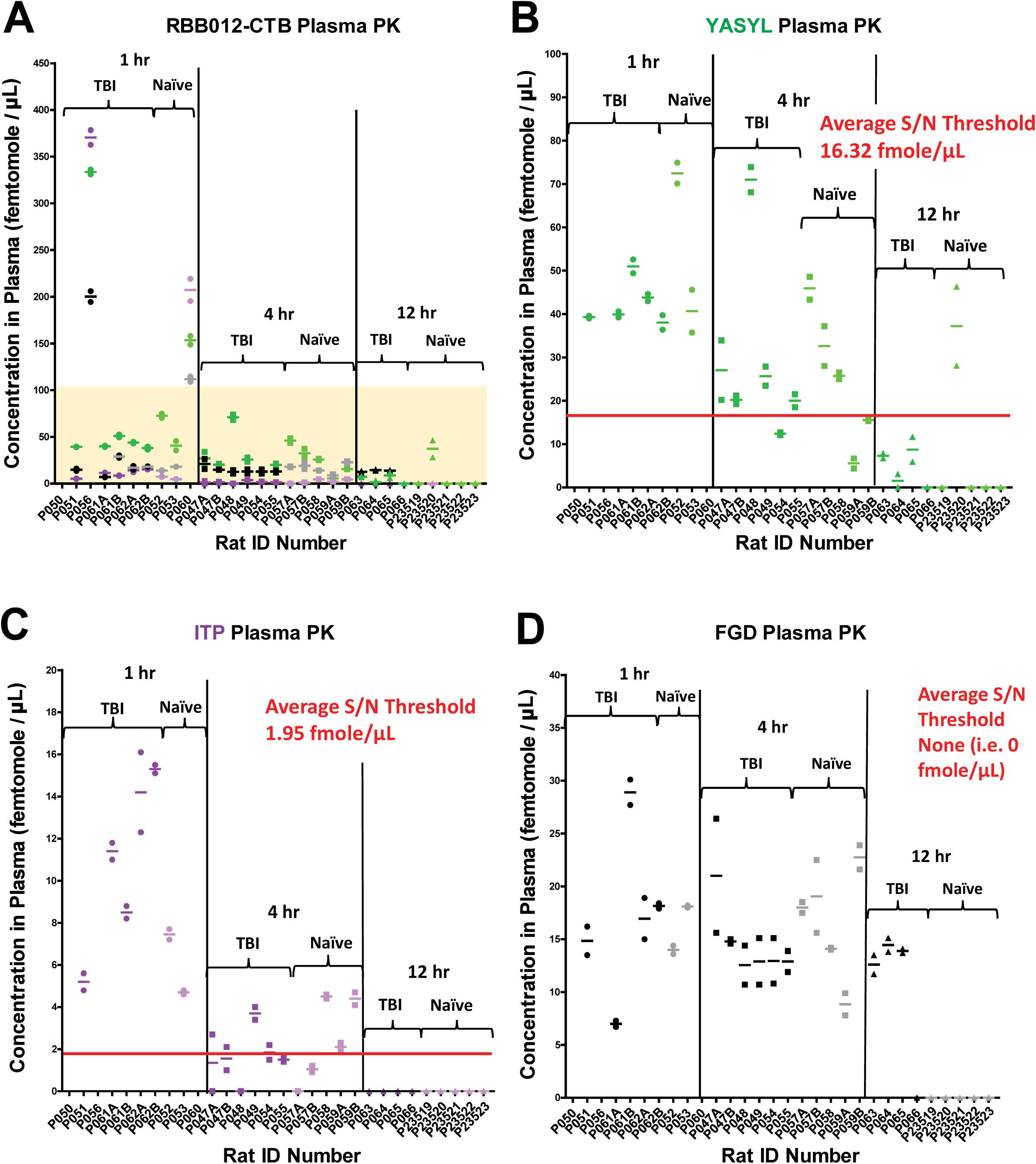
**A)** A composite graph of the 3 peptide markers for RBB012-CTB helps reveal similarities in plasma concentration in the blood. Concentration of the peptide markers is plotted for each rat (whose ID numbers comprise the x-axis). Scatter plots are color coded according to the schematic in Figure 8. Darker colors represent the plasma concentrations determined from rats having incurred a cortical impact (TBI). Pastel colors represent the corresponding peptide concentrations from uninjured rats (“Naïve”). Region highlighted in yellow is the focus of the remaining panels. **B)** A scatter plot showing only the YASYL fragment helps track the variable light chain of the 3E10 subdomain comprising RBB012-CTB. The 2 technical repeats are plotted for each rat along with the average represented as a dashed line. An average signal:noise (S/N) threshold (i.e. the concentration most likely to comprise the LLOQ) is represented by a red line. In this case, the LLOQ is 16.32 fmole/μL. **C)** Scatter plot for the ITP peptide tracking the HSP72 domain of RBB012-CTB. **D)** Scatter plot for the FGD peptide tracking the HSP72 domain and its relative exposure to oxidation.

The LLOQ is different for each peptide given their inherent biochemical differences. For instance, one peptide may fly through the mass spectrometer better than others. Two of the peptides (FGD and YASYL) are more polar than the other peptides investigated; hence, they may not stick to the hydrophobic C18 column similar to other peptides prior to injection into the mass spec. There was no significant difference between the plasma concentrations in TBI injured or naïve rats based on a Welch’s t-test of both groups at each time point. Nor was there a difference in the rate of clearance based on a statistical analysis of linear regressions that is discussed in the next section. Statistical comparisons could not be conducted on the ITP and FGD plasma results at 12 hours since the naïve rats were at 0 fmoles/µL.

### Calculating a Lower Limit of Quantitation (LLOQ)

The average signal to noise threshold does NOT represent a firm demarcation of all concentration values either having an S/N ratio ≥10:1 or <10:1. We have obtained concentration values below the calculated LLOQ that have ratios ≥10:1 (indicating a high confidence in the result) and, conversely, we have obtained concentration values much greater than the LLOQ but a low confidence of being accurate (due to S/N <10:1).

To calculate an average signal to noise threshold, we have taken the values below an S/N ratio of 10:1 and added them up, then divided by the number of these samples. To be clear, all concentration values from injured and naïve rats below an S/N ratio of 10:1 from the 1, 4 and 12 hr samples are used to determine the average. This provides an average LLOQ value for graphing purposes.

We also believe this gives the reader a sense of the most probable concentration limit for the LLOQ. As seen in **Figure 9**, you will note that a significant minority of concentration values are below the LLOQ. Rather than drop these values from the graphs and give the appearance of missing data, it made more sense to plot everything and show how the plasma concentrations dropped asymptotically toward zero over time.

### Crossing the Blood Brain Barrier

Like the biochemical biomarker studies discussed above, the PK studies for brain tissue uptake required harvesting and analysis of the ipsilateral and contralateral hemispheres (**Figure 10A**). Surprisingly, only ITP was detectable in the brain tissue above the LLOQ and it performed well for all 3 Fv-HSP72 variants. Scatter and box plots of the RBB012-CTB results illustrate the data collected by injecting ipsi- and contralateral tissue extracts twice into the mass spec for each rat (**Figure 10B,C**). The plots show a wider distribution of normalized drug concentration in the brain 1h post-injection, both in CCI rats and naïve rats only exposed to anesthesia. Distribution of the pg/mg concentrations narrows within 12h for the TBI rats, while the ITP biomarker is undetectable in Naïve ones (**Figure 10C**). For the HSP72 subunit of our drug, the difference in biodistribution between TBI and Naïve rats at 12h are clear. But we wanted to further analyze the rate of clearance from the brain in both groups and see if statistically significant differences could be discerned before the 12h time point.

**Figure 10.**
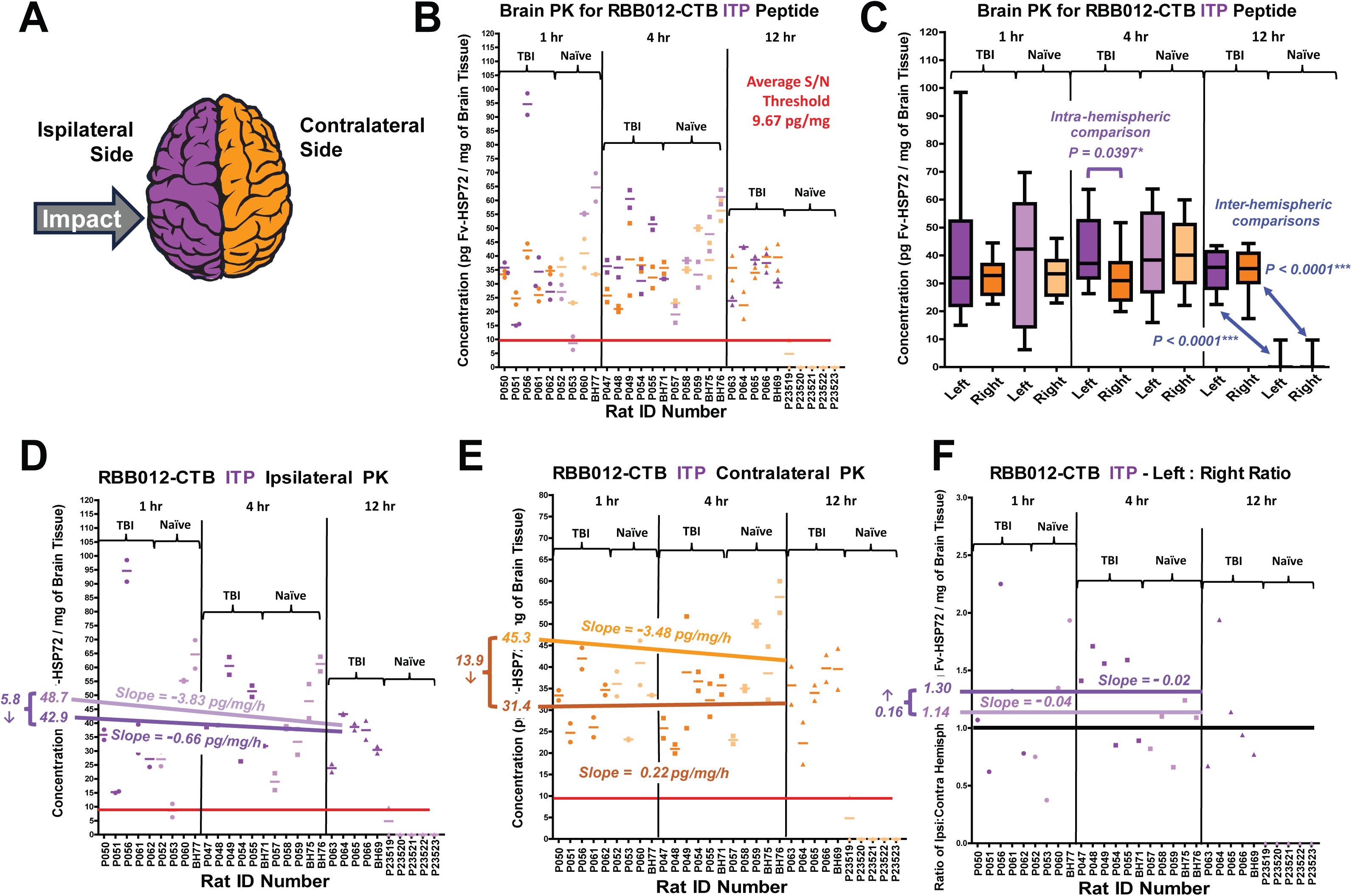
**A)** Schematic showing color coding of the **B)** scatter plots and **C)** box plots comparing RBB012-CTB concentration in both hemispheres of the brain. The 2 technical repeats are plotted for each rat along with the average represented as a dashed line. An average signal:noise (S/N) threshold (i.e. the concentration most likely to comprise the LLOQ) is represented by a red line. Drug concentrations reported in pg of ITP peptide per milligrams of brain tissue. Calculations to normalize the results discussed in Methods. Darker colors (ipsi=purple; contra=tan) represent the normalized concentrations determined from TBI rats. Pastel colors (ipsi=violet; contra=orange) represent the corresponding concentrations from Naïve rats. Distributions are not statistically different for TBI and Naïve rats at 1h post-IV. At 4h, when drug is already reaching unquantifiable levels in plasma, there is a significant difference in distribution between the ipsi- and contralateral hemispheres of the brain (p=0.0397) only in those rats exposed to a CCI. By 12h, the ITP is undetectable in naïve rats and its distribution is similar in both hemispheres of the brain (Inter-hemispheric: p<0.0001). **D)** A linear regression of the ipsilateral ITP concentrations in Naïve rats defines a significant clearance rate of -3.83 pg/mg/hr; however, in TBI rats this rate is changed significantly by 3.17pg/mg/h to a slower clearance rate of -0.66pg/mg/h (-3.83 + 3.17 = -0.66). Projecting both slopes to the y-intercept results in starting concentrations of 48.7 and 42.9pg/mg for the Naïve and TBI rats, an insignificant difference of only 5.8pg/mg. **E)** A similar regression model for the contralateral concentrations determined a significant clearance rate of - 3.48pg/mg/h for Naïve rats and a significant drop in that rate by 3.7pg/mg/h to a rate of 0.22pg/mg/h (-3.48 + 3.7 = 0.22). In this instance, naïve rats have a starting concentration of 45.3pg/mg; however, TBI exposed rats have a significantly lower one at 31.4pg/mg, which is not unexpected given the intra-hemispheric difference seen at 4 hours in (B). **F)** Breaking the intra-hemispheric analysis down further to account for individual rat variability we see statistically similar clearance rates for Naïve and TBI rats (-0.04 vs -0.02) at 1h and 4h. The analysis also confirms greater uptake of drug in the ipsilateral versus contralateral for those rats exposed to a CCI, albeit the 0.16 increase was not statistically significant. Note, these are unitless ratios.

To quantify peptide uptake and clearance over time, our statistician fit linear regression models relating measured peptide concentrations to time, injury group (TBI vs Naïve), and their interaction (i.e., an injury group-specific time trend). The two regression coefficients of each resulting line (y = **m**x + **b**), with **m = the slope** and **b = the y-intercept**, were then used for the statistical analysis. Linear regressions were limited to the 1h and 4h time points. In this parameterization, the “baseline” y-intercept represents the estimated baseline concentration for the Naïve group at time T = 0h, meaning the theoretical amount of ITP, and presumably Fv-HSP72, initially taken up by the brain tissue. The main coefficient for time is the rate of drug clearance in Naïve rats as represented in the slope of the linear regression for that injury group. Shifts in the **b**-coefficient, representing the shift in the T=0h intercept for TBI rats relative to Naïve ones were determined; as were shifts in the **m**-coefficient (slope) for TBI animals versus Naïve ones. We tested whether baseline levels (intercepts) and clearance rates (slopes) differed between TBI and Naïve rats by examining the corresponding regression coefficients and their standard errors from these fitted models. Analyses were performed separately by peptide for each of the Fv-HSP72 variants (although only RBB012-CTB is presented here) in both plasma (**Figure 9**) and brain (**Figure 10**) samples. Brain-tissue models were fitted separately for ipsilateral and contralateral hemispheres.

For the ipsilateral data sets (**Figure 10D**), there is a significant decrease in the RBB012-CTB clearance rate from 1h to 4h in the Naïve rats: -3.83 pg/mg/h (p < 0.01). This clearance rate *changed* significantly by 3.17pg/mg/h (p<0.05) when measured in TBI rats, resulting in a clearance rate of only -0.66pg/mg/h. The theoretical drug uptake at T = 0h was not significantly different for TBI and Naïve rats. Simply put, injury to the brain results in a longer retention time for RBB012-CTB in the ipsilateral hemisphere compared to Naïve rats. For the contralateral data sets (**Figure 10E**), there is also a significant decrease in the clearance rate from 1h to 4h in Naïve rats: -3.48pg/mg/h (p < 0.01). Despite the injury being in the opposite hemisphere of the brain, the clearance rate also *changed* significantly by 3.7pg/mg/h (p < 0.01) in TBI rats yielding a fitted TBI slope near zero (0.22 pg/mg/h). Given the uncertainty in the fitted coefficients, this point estimate should be *<u>interpreted as strong evidence of substantially reduced clearance / increased</u> <u>retention in CCI injured rats,</u>* rather than definitive evidence of net accumulation. Additionally, the fitted intercept is lower by 13.9 pg/mg (p < 0.05). One possible explanation for such a decrease could be redistribution toward the ipsilateral injury site, a hypothesis supported by the significant hemispheric localization difference at 4h in CCI-injured rats (**Figure 10C**, p = 0.0397). Analysis of intra-hemispheric differences in individual rats at 1h and 4h revealed statistically similar accumulation and clearance of the Fv-HSP72 in both hemispheres of the brain regardless if the animal was exposed to a TBI or not; albeit there is a small, but statistically insignificant (1.30 vs 1.14), increase in drug uptake in the ipsilateral side of CCI injured rats (**Figure 10F**). Changes in clearance rates must be significant after the first 4 hours to account for the biodistribution differences seen at 12h.

## Discussion

A large body of evidence demonstrates HSP72 as one of the body’s therapeutic responses to tissue damage regardless of the cause of the trauma. HSP72 is a pleiotropic agent originally shown to refold denatured proteins in order to prevent intracellular protein aggregation^25^. It also inhibits apoptosis that occurs through at least three different pathways (the ATP-dependent apoptosome, ATP-independent AIF and the NF-κB pathways)^26–28^. The NF-κB pathway is particularly interesting since it is triggered in the brain’s vascular system at the penumbra of a contusion^29^. In a separate study, HSP72 upregulation in cerebral microvascular endothelial cells occurred, via cytokine FGF21, in response to hypoxic stress^30^. Fv-HSP72 may have the potential of inhibiting delayed vascular destruction in the brain post-CCI. Until now, the problem has been the inability to *rapidly* deliver consequential amounts of HSP72 into cells in damaged tissues. The strategy of fusing a cell penetrating antibody fragment (mAb 3E10), that specifically targets areas of cellular destruction, with HSP72 results in a biologic (Fv-HSP72) that can work as an effective cytoprotectant *<u>after</u>* traumatic injury. Some propose using small molecules to induce this key cytoprotectant^31^, but such methods of HSP72 induction have a lag time and heat shock protein induction attenuates with aging^32–35^. Rather than wait hours for induction, Fv-HSP72 rapidly places exogenous HSP72 to the area of traumatic injury, eliminating any lag time, both in young and old.

Expanding on work by others showing HSP72 inhibiting brain lesions in mouse cortical impact models, we screened three Fv-HSP72 structural variants (**Figure 1**) engineered by Rubicon Biotechnology in a rat CCI model simulating a moderate to severe impact. Ipsilateral and contralateral brain tissue extracts were probed for biomarkers of neurodegeneration (pTau, NF-L, pNF-H), oxidative stress (3-NT) and inflammation (IL-6). Two variants (RBB010 and RBB012) possessed a Swivel linker and a third possessed a proprietary cathepsin B cleavable linker (RBB012-CTB). All three were tested at two dosages 10 and 30mg/kg. RBB012-CTB proved to be superior in reducing biomarker levels and is now in development as a TBI therapeutic.

In controlled cortical impact studies of TBI, it is important to compare the results in the ipsilateral (left) hemisphere with the contralateral (right) in each rat to reduce the level of variability across a population of animals. Our results demonstrate it is important to statistically analyze ***intra***-hemispheric and ***inter***-group differences. As seen with inter-group analyses of biomarkers pTau S396, 3-NT and IL-6, treatment with RBB012-CTB dropped absorbance readings to levels comparable with naïve rats (**Figures 1C,6B,7D**); yet, these differences are not fully reflected by only evaluating intra-hemispheric left:right brain ratios (**Figures 1D,6C,7F**). Five biomarkers of neurodegeneration were analyzed, and while RBB012-CTB decreased absorbances for the pTau S396 biomarker in both hemispheres of treated rats (**Figure 1**), other markers (pTau T231, T181 and NF-L) exhibited hemispheric localization with biomarker decreases in the ipsilateral extracts for rats receiving drug versus those given vehicle buffer (**Figures 2, 3 and 4**). The pNF-H levels were dose-dependent, only showing a significant decrease at 30mg/kg (**Figure 5**). Results suggest RBB012-CTB localizes to the injured tissues in the ipsilateral hemisphere, where extracellular DNA would be present, reducing neurodegeneration in the area of the brain that was impacted. Silver staining in rodent models has demonstrated the contralateral cortex can be injured over time due to anterograde degeneration of projections from the ipsilateral hemisphere^17^. Future studies will evaluate the benefits of increased and repeat dosing post-injury to reduce biomarker levels, and provide sufficient Fv-HSP72 protection, to both hemispheres. In certain circumstances, the surgical procedure alone increased neurodegeneration biomarker levels in sham rats (see **Figures 1C** and **5D** for pTau S396 and pNF-H, respectively); but treatment with RBB012-CTB resulted in absorbances below even the sham controls. Even with great care taken during surgery, some neural tissue damage is to be expected; however, this highlights Fv-HSP72’s potential broader clinical use, including reducing degeneration after neurosurgery, or in less severe forms of TBI.

Fv-HSP72 was originally developed to reduce damage caused by oxidative stress when the occluded blood vessels of neural and cardiac tissues are reperfused^6,14^. Hence, it was reasonable to investigate whether the drug could inhibit nitration of tyrosine residues by RNS, a known biomarker of radical activity in TBI models. Significant reductions were seen in both hemispheres of the brain with either dosage of RBB012-CTB (**Figure 6**). We also evaluated IL-6, a subacute inflammatory biomarker whose levels increase in the days and weeks post-injury^16^. At 30mg/kg, IL-6 levels were indistinguishable from naïve rats (**Figure 7**).

Disruption of the BBB after a direct impact has been well documented^36–38^, with 70% of TBI patients in one clinical study exhibiting BBB breakdown as determined by MRI and EEG measurements^36^. We investigated multiple tryptic peptide sequences unique to each of the Fv-HSP72 variants and identified ones suitable for tracking and quantifying blood clearance and brain uptake using MS (**Figure 8**). When analyzing plasma levels, these peptide biomarkers approached the LLOQ for MS detection after 4 hours (**Figure 9**). A similar clearance pattern was seen for all three MS peptide biomarkers for both RBB012 and RBB012-CTB (data not shown).

A subset of samples, particularly at the 12h time point, were undetectable by MS, even if they were detected in the HPLC prior to injection into the spectrometer. Increasing the injection volumes for these particular samples may have resulted in detectable signals, but there was no point doing that when the quantities are so low and below the LLOQ. Thus, we decided that these sample that were *<u>undetectable by MS should be given the designation of ZERO fmol/μL by convention</u>*. Furthermore, since many samples at 4 and 12 hrs were assigned 0 fmol/μL, we decided to not count these 0 fmol/µL samples in the LLOQ value in order not to artificially “deflate” the LLOQ to near zero. As already mentioned, there are samples in the data set with high concentration values but a low confidence of being accurate (due to S/N <10:1). A further deflation of the LLOQ can result in a greater number of “unreliable” concentrations values with S/N ratios <10:1 appearing above the LLOQ limit. We want to give the reader a sense of the most probable concentration where the LLOQ is located.

In brain tissue, only the ITP peptide was detectable above the LLOQ by MS (**Figure 10**). Having no *unoxidized* FGD detected above its LLOQ suggests the drug was exposed to significant oxidation at all time points once it entered the brain whether the rat was subjected to a CCI or not. Toward the N-terminus of RBB012-CTB, we track the 3E10 scFv subdomain using YASYL. Lack of YASYL detection may be related to DNA binding and not necessarily side-chain oxidation. If that is the case, it will be difficult to determine a specific range of molecular weights to screen for using MS given the wide diversity of sizes we would have to search if the 3E10 subunit is binding random fragments of DNA. There is a second plausible explanation. While conducting our experiments, a study was published by another research group demonstrating that the 3E10 antibody fragment that binds DNA, once it enters a cell, is transported to the nucleus by the protease cathepsin B and degraded^39^. Since the Fv-HSP72 has protease cleavable linkers, it is entirely possible that once the fusion protein rapidly enters brain tissue, the 3E10 antibody subunit is cleaved and degraded by cathepsin B within the nucleus; hence, escaping detection.

A caveat in the MS analyses is that the majority of mass spec values at 12h, in particular for naïve rats, were below the LLOQ. Since values below the LLOQ were considered “unreliable”, they were treated as missing values by the statistician and these covariates could not be included in the statistical models. *<u>Importantly, this led to an overly conservative statistical comparison</u>* of the TBI and Naïve groups given the 12h results could not be compared (even if it is clear from the data that there is a significant difference in ITP peptide marker between the groups of rats).

While the clearance of our Fv-HSP72 from the brains of Naïve rats at 12h was obvious, linear regression models were made of the 1h and 4h data sets and the variances subjected to statistical analysis. In both hemispheres, the slopes were so small they effectively represent a very slow clearance of RBB012-CTB in those rats suffering a CCI injury (**Figures 10D,E**). Note, the theoretical protein uptake into the brain at T=0h for TBI and Naïve rats in both hemispheres was 42.9, 45.3 or 48.7pg/mg, and yet, the contralateral TBI rats drop to 31.4pg/mg; this significant difference in drug detection may be attributed to greater RBB012-CTB accumulation in the ipsilateral tissues of the injured animals. One question that must be addressed is the accumulation of our Fv-HSP72 in the brains of Naïve rats. This raises the question of BBB disruption in anesthetized animals. Our Naïve rats were anesthetized with isoflurane and this may have been sufficient to temporarily disrupt the BBB as researched by one team that found permeability of 10kD dextran post-anesthesia in mice; but, notably, they did not see permeability of 70kD dextran^40^. BBB disruptions are often temporary, with one CCI study observing exogenous horseradish peroxidase uptake in the ipsilateral and, to a lesser extent, contralateral hemispheres up to 3 hours post-injury^41^. Another study observed the influx of liposomes up to 8 hours post-injury^38^. While this observation requires further study, it is clear that between 4 and 12 hours, RBB012-CTB washes out of the brain tissues of naïve rats while still detectable in both hemispheres of TBI animals. Analyzing the ratio of ipsi-versus contra-distributions in the TBI rats reveals variation in drug retention between the two halves dissipates by 12h (**Figure 10F**), possibly due to the Fv-HSP72 targeting further cell death as it spreads via axonal degeneration across hemispheres^17,42^.

Key to the uptake of 98kD Fv-HSP72 fusion protein is the ENT2 channel, through which molecules can pass based on equilibration, not requiring the energy required of endocytosis and transporters. ENT2 gene expression occurs in many cell types, including brain and vascular cells^9^; the specificity of the Fv-HSP72 therapeutic is the targeting of tissues undergoing significant damage where extracellular DNA is abundant and easily accessible. Salvaging of DNA by surrounding cells through the ENT2 channel provides 3E10^7,8^, and any attached payloads, the ability to enter those cells still viable and rescue them. Thus, the Fv-HSP72 cytoprotectant is capable of rescuing multiple tissue types, as evidenced by statistically significant *in vivo* results in brain tissue with a rat stroke model^6^, cardiac tissue with a rabbit myocardial infarction model^14^, lung tissue with a rat phosgene intoxication study^43^ and retinal tissue in a rat blast overpressure model (manuscript in preparation).

In summary, our strategy of rapid HSP72 delivery to prevent symptoms leading to axonal degeneration, oxidative stress and neuronal cell death can improve on nature’s own countermeasures to combat cell loss. Specifically, in a focal contusion model of TBI, rapid HSP72 delivery to brain tissue reduced neurodegeneration, oxidative stress and inflammation. Importantly, the RBB012-CTB variant of Fv-HSP72 has great potential as a TBI therapeutic.

## Transparency, Rigor and Reproducibility Summary

Key aspects of the experimentation and analysis were delegated to Rubicon, JHU and Stanford to minimize risk of bias during the study. Rubicon synthesized the drugs and tested their efficacy *in vitro* prior to transfer to JHU for animal testing. All rats were randomized and provided ID numbers by JHU and the blinded tissue samples for the biomarker studies were sent to Rubicon, while blinded samples for the mass spec studies were sent to Stanford. Biomarker and mass spec data were unblinded by Parseghian prior to plotting the results. For the biomarker studies, we performed a power analysis to determine a sample size, based upon parameter estimates of standard deviations and effect size from prior work in a focal TBI model^20^, that would allow us to see, with 80% power and a family-wise type I error rate of 0.05 (after Bonferroni correction, α=0.017 per comparison) a 25% difference in the primary endpoint between vehicle and treated groups in an experiment in which we would simultaneously compare vehicle to each of 3 drug doses. Based upon that analysis, we chose an N of 8 rats per group. The number of rats per group were chosen based on an assumption of a 20-25% loss due to experimental complications.

## Author’s Contributions

Investigation (tissue extractions & biomarker analysis): AC, KP, SS, AS; Investigation (CCI model & tissue extractions): MS, CLP, PA; Investigation (mass spec): KK; Conceptualization & Methodology: DH, RN, SH, KK, CLP, JRO, PA, MHP; Data Analysis (biomarker & mass spec): DH, RN, KK, CLP, MHP; Data Curation: MHP; Formal Analysis: PP, JRO, MHP; Resources (synthesis and purification of Fv-HSP72): RAR, AC; Project Administration: RAR, CLP, MHP; Funding Acquisition: RAR, MHP; Writing & Visualization: MHP; Review & Editing: All authors except SS and AS.

## Disclaimer

The views expressed in this manuscript reflect the results of research conducted by the author(s) and do not necessarily reflect the official policy or position of the Defense Health Agency, Department of War, nor the U.S. Government.

## Acknowledgements

We thank Grace Chan at Rubicon and Jacquelyn Saunders at the UCLA/Vet Affairs lab for their excellent technical assistance, as well as Soren Faulkner for outstanding assistance running the hundreds of MS samples at Stanford. We also thank Sarah Fontaine, PhD and Colleen P. Lavinka, PhD of the CDMRP for helpful technical discussions. This work was partially supported through the Congressionally Directed Medical Research Programs by the Office of the Assistant Secretary of Defense for Health Affairs through the Defense Medical Research and Development Program and endorsed by the War Department **Award No. W81XWH-19-2-0027**. Opinions, interpretations, conclusions and recommendations are those of the author and are not necessarily endorsed by the War Department.

## Abbreviations

CHO: Chinese hamster ovary
DTT: Dithiothreitol
ENT2: equilibrative nucleoside transporter 2
HSP: heat shock proteins
h or hr(s): hour(s)
IVT: intravitreal
IV: intravenous
msec: milliseconds
m: minutes
mAb: monoclonal antibody
s: seconds
TBS: Tris Buffered Saline
TUNEL: terminal deoxynucleotidyl transferase dUTP nick end labeling
TBI: traumatic brain injury

